# The catalase contributes to microaerophilic H_2_O_2_ priming and the peroxiredoxins AhpC, Tpx and Bcp confer resistance to organic hydroperoxides in *Staphylococcus aureus*

**DOI:** 10.1101/2022.07.19.500589

**Authors:** Nico Linzner, Vu Van Loi, Haike Antelmann

**Author notes:** **Corresponding author:** Haike Antelmann, Institute for Biology-Microbiology, Freie Universität Berlin, Königin-Luise-Straße 12-16, D-14195 Berlin, Germany.

## Abstract

*Staphylococcus aureus* is a major human pathogen, which has to cope with oxidative stress as part of the host innate immune defense under macrophage and neutrophil infections. In this study, we have investigated the role of the catalase KatA and the peroxiredoxins AhpC, Tpx, and Bcp for priming and resistance under oxidative stress in *S. aureus* during aerobic and microaerophilic growth. The results revealed that *S. aureus* is resistant to high doses of up-to 100 mM H_2_O_2_ during the aerobic growth. While KatA is essential for this high aerobic H_2_O_2_ resistance, the peroxiredoxin AhpC contributes to detoxification of 0.4 mM H_2_O_2_ in the absence of KatA. In addition, AhpC, Tpx and Bcp were shown to be required for detoxification of cumene hydroperoxide (CHP) and regeneration of the reduced state of the bacillithiol (BSH) redox potential during recovery from CHP stress in *S. aureus*. The high H_2_O_2_ tolerance of aerobic *S. aureus* cells was associated with priming by endogenous H_2_O_2_ levels, which was supported by an oxidative shift of the basal level of *E*_BSH_ (−291 mV) compared to that in microaerophilic cells (−310 mV). In contrast, *S. aureus* can be primed by sub-lethal doses of 100 µM H_2_O_2_ during the microaerophilic growth to acquire an improved resistance towards the otherwise lethal triggering stimulus of 10 mM H_2_O_2_. This microaerophilic priming was dependent on increased *katA* transcription and elevated KatA activity, whereas aerobic control cells showed already constitutive high KatA activity. Thus, KatA is the major player contributing to the high H_2_O_2_ resistance of aerobic cells and to microaerophilic H_2_O_2_ priming to survive the subsequent lethal triggering doses of H_2_O_2_, allowing the adaptation of *S. aureus* to oxidative stress under infections in different oxygen environments.

## INTRODUCTION

*Staphylococcus aureus* is a major human pathogen, which can cause local skin and soft tissue infections, but also live-threatening diseases, such as septicaemia, endocarditis, necrotizing pneumonia and osteomyelitis (Archer, 1998;Lowy, 1998;Boucher and Corey, 2008). During infections, *S. aureus* has to cope with the oxidative burst of professional phagocytes, such as macrophages and neutrophils (Linzner et al., 2021). The NADPH oxidase in the phagosomal membrane generates superoxide anion (O_2_•^-^), which is converted to H_2_O_2_ by the superoxide dismutase (Winterbourn and Kettle, 2013;Winterbourn et al., 2016). The myeloperoxidase MPO further converts H_2_O_2_ with chloride to the highly toxic HOCl, which is released to kill the invading pathogens (Ulfig and Leichert, 2021). In addition,

*S. aureus* encounters reactive oxygen species (ROS) during incomplete reduction of molecular oxygen in the respiratory chain (Imlay, 2003). ROS can damage all cellular macromolecules, resulting in oxidation of proteins, lipids and carbohydrates (Imlay, 2008;Linzner et al., 2021). *S. aureus* uses various enzymatic and non-enzymatic ROS detoxification systems, such as the catalase (KatA), peroxiredoxins (AhpC, Tpx, Bcp) and the low molecular weight (LMW) thiol bacillithiol (BSH) (Linzner et al., 2019;Linzner et al., 2021). BSH is associated with the bacilliredoxin/BSH/YpdA redox pathway to regenerate oxidized protein thiols and bacillithiol disulfide (BSSB) (Linzner et al., 2019). We have previously constructed a Brx-roGFP2 fused biosensor to monitor the changes in the BSH redox potential (*E*_BSH_) under oxidative stress in *S. aureus* (Loi et al., 2017). This study already revealed that *S. aureus* is highly resistant to 100 mM H_2_O_2_, since the Brx-roGFP2 biosensor responded only weakly to high H_2_O_2_ levels, leading to small *E*_BSH_ changes (Loi et al., 2017). The catalase KatA was identified as the major H_2_O_2_ detoxification enzyme, which conferred the constitutive H_2_O_2_ resistant phenotype to aerobically grown *S. aureus* cells (Horsburgh et al., 2001;Weber et al., 2004). KatA is also important for nasal colonization and mediates protection under macrophage and neutrophil infections (Mandell, 1975;Horsburgh et al., 2001;Cosgrove et al., 2007;Das et al., 2008). The peroxiredoxin AhpCF showed compensatory roles in resistance to H_2_O_2_ and organic hydroperoxides (OHPs) and contributed to nasal colonization (Cosgrove et al., 2007).

In generally, AhpC, Tpx and Bcp can be classified into typical (AhpC) and atypical (Tpx, Bcp) 2-Cys peroxiredoxins based on their thiol-oxidation mechanism between the peroxidatic (C_P_) and resolving Cys (C_R_), involving inter- or intramolecular disulfides, respectively (Poole et al., 2011). The functions and substrates of AhpC, Tpx and Bcp have been previously studied in *Escherichia coli*. AhpC detoxification of H_2_O_2_ leads to formation of an oxidized AhpC dimer, which aggregates to an oligomer with chaperone functions (Poole et al., 2011;Parsonage et al., 2015). Regeneration of AhpC requires the NADPH-dependent flavin disulfide reductase AhpF as redox partner (Poole et al., 2011;Parsonage et al., 2015). The peroxidase Bcp of *E. coli* is induced by H_2_O_2_, OHPs and during aerobic growth and conferred resistance against H_2_O_2_ and OHP stress (Jeong et al., 2000). The thiol-peroxidase Tpx of *E. coli* was shown to catalyze detoxification of H_2_O_2_ and OHPs *in vitro* and is recycled by the Trx/TrxR system (Baker and Poole, 2003;Cha et al., 2004). In *S. aureus*, Tpx responds strongly to H_2_O_2_ and other thiol-reactive compounds and was oxidized in the redox proteome under HOCl stress (Imber et al., 2018;Linzner et al., 2021). However, the detailed functions of the peroxiredoxins AhpC, Tpx and Bcp in peroxide resistance, detoxification and survival have not been studied thus far in *S. aureus*.

In *S. aureus*, transcription of *katA, ahpCF* and *bcp* is strongly induced only by high levels of 10 mM H_2_O_2_ and controlled by the peroxide-responsive PerR repressor (Horsburgh et al., 2001;Chelikani et al., 2004;Cosgrove et al., 2007). In *Bacillus subtilis*, KatA is also a member of the PerR regulon, but already inducible by sub-lethal doses of 100 µM H_2_O_2_ (Mongkolsuk and Helmann, 2002;Faulkner and Helmann, 2011). Pretreatment of *B. subtilis* cells with sub-lethal H_2_O_2_ as “priming stimulus” confers improved resistance towards subsequent lethal H_2_O_2_ doses, termed “triggering stimulus”, which is encountered as future stress (Murphy et al., 1987;Engelmann and Hecker, 1996). This H_2_O_2_ priming effect was shown to be mediated by KatA, which is induced by the mild stress to prepare the cells for better survival of lethal future oxidative stress (Engelmann and Hecker, 1996). Similarly, H_2_O_2_ priming for improved resistance towards the triggering stimulus was dependent on the OxyR-dependent enzymes KatG and AhpCF in *E. coli* and *Salmonella* Typhimurium (Christman et al., 1985;Faulkner and Helmann, 2011;Chiang and Schellhorn, 2012). Although *S. aureus* exhibits a constitutive H_2_O_2_ resistance during the aerobic growth, it is unknown whether priming for improved H_2_O_2_ resistance is possible under aerobic or microaerophilic conditions.

In this study, we used growth and survival phenotype analyses, Brx-roGFP2 biosensor measurements and transcriptional studies to investigate the functions of KatA and the peroxiredoxins AhpC, Tpx and Bcp in peroxide resistance, detoxification and priming during aerobic and microaerophilic growth. Our results showed that *S. aureus* is H_2_O_2_ primable for improved resistance only under microaerophilic conditions, which is dependent on KatA. In contrast, aerobic growth already leads to increased levels of ROS, which causes KatA-dependent aerobic priming for constitutive H_2_O_2_ resistance. While KatA confers H_2_O_2_ resistance in *S. aureus*, the peroxiredoxins AhpC, Tpx and Bcp were shown to contribute to survival and resistance under CHP stress and regeneration of reduced *E*_BSH_ upon recovery from CHP stress.

## MATERIALS AND METHODS

### Bacterial strains, growth and survival assays

Bacterial strains, plasmids and primers are described in **Table S1-S3**. For genetic manipulation, *E. coli* was cultivated in Luria Broth (LB) medium. *S. aureus* COL strains were grown in RPMI medium to an optical density at 500 nm (OD_500_) of 0.5 and exposed to H_2_O_2_, cumene hydroperoxide (CHP) or hypochlorous acid (HOCl), followed by determination of colony forming units (CFUs) in survival assays as described (Loi et al., 2018). Each experiment was performed in at least 3 independent biological replicates and the results are presented as mean values with standard deviations (SD) from all biological replicates as indicated in each figure legends. Statistical analysis was performed using the Student’s unpaired two-tailed *t*-test by the graph prism software. The biochemical compounds were purchased from Sigma Aldrich. The HOCl concentration was determined as described (Winter et al., 2008).

### Construction of the *S. aureus* COL Δ*katA*, Δ*ahpC*, Δ*ahpC*Δ*katA*, Δ*tpx*, Δ*bcp* and *perR* mutants and complemented strains

The *S. aureus* COL Δ*katA* mutant and *katA* complemented strains were previously constructed (Linzner et al., 2020). The *S. aureus* Δ*ahpC*, Δ*tpx*, Δ*bcp* and Δ*perR* deletion mutants were constructed using the temperature-sensitive *E. coli-S. aureus* shuttle vector pMAD as described (Arnaud et al., 2004). In brief, 500 bp of the up- and downstream flanking regions of the specific genes were fused by PCR, digested with *Bgl*II and *Sal*I and ligated into pMAD. The constructs were electroporated into the restriction-negative *S. aureus* RN4220, followed by phage transduction using phage 81 into *S. aureus* COL (Rosenblum and Tyrone, 1964). For construction of the Δ*ahpC*Δ*katA* double mutant, the plasmid pMAD-Δ*katA* of *S. aureus* RN4220-pMAD-Δ*katA* was transduced by the phage 81 into *S. aureus* COL Δ*ahpC* mutant. Selection of the Δ*ahpC*, Δ*bcp*, Δ*tpx*, Δ*perR* and Δ*ahpC*Δ*katA* deletion mutants was performed as described (Loi et al., 2018).

Construction of the His-tagged *S. aureus ahpC, bcp* and *tpx* complemented strains was performed using the plasmid pRB473 as described (Loi et al., 2017). The genes were cloned into pRB473 after digestion with *Bam*HI and *Kpn*I/ *Sac*I, resulting in plasmids pRB473-*ahpC-His*, pRB473-*bcp-His* and pRB473-*tpx-His*, which were transduced in the Δ*ahpC*, Δ*bcp* and Δ*tpx* deletion mutants. In addition, the plasmid pRB473-*brx-roGFP2* (Loi et al., 2017) was introduced into the *S. aureus* COL Δ*katA*, Δ*ahpC*, Δ*ahpC*Δ*katA*, Δ*tpx* and Δ*bcp* deletion mutants to construct the Brx-roGFP2 biosensor expressing catalase and peroxiredoxin-deficient mutant strains.

### Priming and triggering experiments

For priming and triggering, the *S. aureus* strains were grown aerobic under vigorous agitation in a shaking water bath at 150 rpm or microaerophilic in 50 ml Falcon tubes including 40 ml cultures with closed lids without shaking as in previous publications (Fritsch et al., 2019;Fritsch et al., 2020). At an OD_500_ of 0.3, naïve *S. aureus* cells were primed with sub-lethal doses of 0.1 or 1 mM H_2_O_2_, respectively, for ∼30 min. Subsequently, the primed cells were exposed to 10 or 40 mM H_2_O_2_, respectively, as lethal triggering stimulus followed by counting of CFUs.

### Brx-roGFP2 biosensor measurement

To monitor the *E*_BSH_ changes by H_2_O_2_ and CHP, we used the Brx-roGFP2 biosensor expressing WT, Δ*katA*, Δ*ahpC*, Δ*ahpC*Δ*katA*, Δ*tpx* and Δ*bcp* mutant strains and performed injection assays with the oxidants. For measurements of Brx-roGFP2 oxidation during microaerophilic and aerobic H_2_O_2_ priming and triggering experiments, the *S. aureus* COL strain expressing Brx-roGFP2 was cultivated in LB medium to an OD_540_ of 0.3 and challenged with the priming dose of 0.1 mM H_2_O_2_ for 30 min, followed by the triggering dose of 10 mM H_2_O_2_ as described above. Samples were harvested at C, P, PT and T, alkylated with 10 mM NEM, washed and resuspended in PBS with 10 mM NEM. The Brx-roGFP2 oxidation degree (OxD) and *E*_BSH_ changes were determined in the *S. aureus* strains during oxidant injection or in samples harvested at C, P, PT and T as previously described (Loi et al., 2017;Loi and Antelmann, 2020). For fully reduced and oxidized controls, biosensor strains were treated with 10 mM DTT and 5 mM diamide, respectively. The Brx-roGFP2 fluorescence emission was measured at 510□nm after excitation at 405 and 488□nm using the CLARIOstar microplate reader (BMG Labtech). The OxD of the Brx-roGFP2 biosensor was determined for each sample and normalized to fully reduced and oxidized controls as described (Loi et al., 2017;Loi and Antelmann, 2020).

### Northern Blot analyses

Northern blot analyses were conducted with RNA isolated from *S. aureus* COL WT cells in the naïve (C), primed (P), primed and triggered (PT) and triggered only (T) state as described (Wetzstein et al., 1992). RNA isolation was performed using the acid phenol extraction protocol. Hybridizations were conducted using digoxigenin-labeled antisense RNA probes for *katA, ahpC* and *dps* that were synthesized *in vitro* using T7 RNA polymerase and the corresponding primers katA-NB-for/rev and ahpC-NB-for/rev as described (Wetzstein et al., 1992). The *dps* antisense RNA probe was constructed previously (Fritsch et al., 2020).

### Determination of the catalase activity using native PAGE and diaminobenzidine staining

To analyze catalase activities in *S. aureus* COL strains, protein extracts were prepared under native conditions and 50 µg of each sample were separated by non-denaturing 10 % polyacrylamide gel electrophoresis. The gel was stained for catalase activity using 50 µg/ml horseradish peroxidase coupled with 5 mM H_2_O_2_ and 0.5 mg/ml diaminobenzidine as described previously (Clare et al., 1984;Loewen and Switala, 1987).

## RESULTS

### *S. aureus* exhibits a KatA-dependent H_2_O_2_ resistance during the aerobic growth

To investigate the roles of the catalases and peroxiredoxins in the constitutive H_2_O_2_ resistance of *S. aureus* COL, phenotype analyses of the Δ*katA*, Δ*ahpC*, Δ*ahpC*Δ*katA*, Δ*tpx* and Δ*bcp* mutants and the complemented strains were performed during the aerobic growth under H_2_O_2_ stress **(Fig 1; Fig. S1 and S2)**. In agreement with previous findings (Cosgrove et al., 2007), the Δ*katA* mutant was strongly impaired in growth after exposure to 10 mM H_2_O_2_ and did not survive doses of 40 mM H_2_O_2_ **(Fig. 1A,E)**. While the *S. aureus* COL wild type (WT) was not impaired in growth by low doses of 1 mM H_2_O_2_ and survived to 330 and 725 % after 2 and 4h, respectively, the Δ*katA* mutant was hypersensitive to peroxide stress, since the growth was inhibited by 0.4 and 1 mM H_2_O_2_ and only 32% and 4% of cells survived the 1 mM H_2_O_2_ treatment after 2 and 4h, respectively **(Fig. 1F; Fig. S1B)**. However, complementation of the Δ*katA* mutant with pRB473-encoded *katA* could only partially restore the H_2_O_2_ resistance to WT level after treatment with 0.4-1 mM H_2_O_2_ **(Fig. 1F; Fig. S1A-C)**. This incomplete recovery of the WT resistance was due to lower catalase activity in the *katA* complemented strain as confirmed using the diaminobenzidine gel staining method **(Fig. S1D)**.

**Fig. 1.**
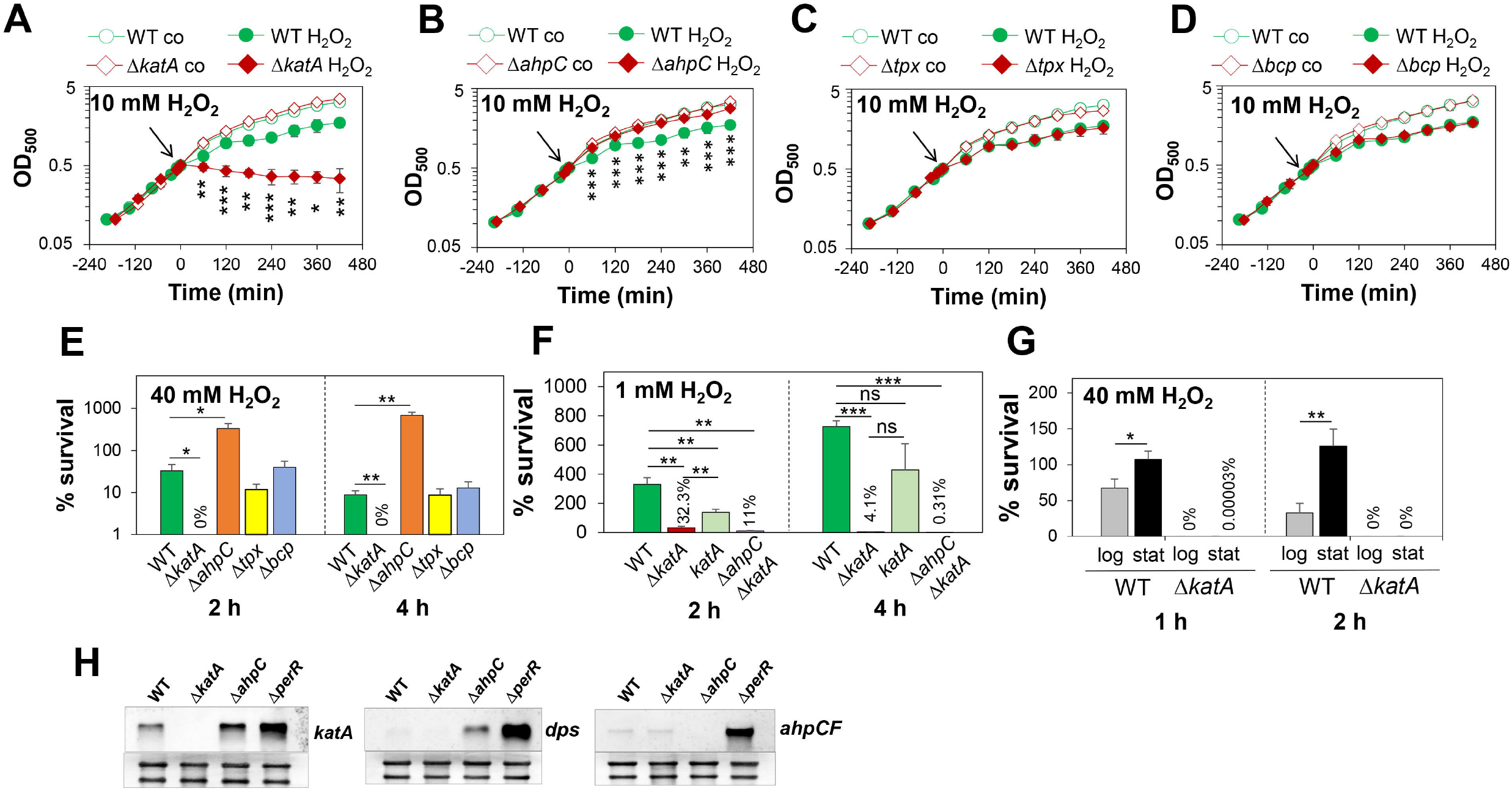
KatA confers the strong H_2_O_2_ resistance during the aerobic growth, but peroxiredoxins are not required for growth and survival under H_2_O_2_ stress. **(A-D)** Growth curves of *S. aureus* COL WT, the Δ*katA* **(A)**, Δ*ahpC* **(B)**, Δ*tpx* **(C)** and Δ*bcp* mutants **(D)** in RPMI medium before (co) after exposure to 10 mM H_2_O_2_ at an OD_500_ of 0.5. **(E)** Survival rates were determined as CFU counts for *S. aureus* COL WT, Δ*katA*, Δ*ahpC*, Δ*tpx* and Δ*bcp* mutants at 2 and 4 h after treatment with 40 mM H_2_O_2_. **(F)** The Δ*katA* mutant is sensitive and the Δ*ahpC*Δ*katA* mutant is hypersensitive to 1 mM H_2_O_2_ as revealed by survival rates based on CFUs counts. **(G)** *S. aureus* COL WT and Δ*katA* mutant cells were exposed to 40 mM H_2_O_2_ during the log and stationary phases at OD_500_ of 0.5 and 2-3, respectively. The survival rates were calculated relative to the untreated control, which was set to 100%. Mean values and standard deviation (SD) of 3-4 biological replicates are shown. The statistics was performed using the Student’s unpaired two-tailed t-test by graph prism. Symbols are: ^ns^p > 0.05, *p ≤ 0.05, **p ≤ 0.01 and ***p ≤ 0.001. **(H)** Northern blot analyses of the *katA, ahpCF* and *dps* specific transcripts of the PerR-regulon in the *S. aureus* WT, Δ*katA*, Δ*ahpC*, and Δ*perR* mutants indicates that the Δ*ahpC* mutant has increased transcript levels of *katA* and *dps* during the log phase at an OD_500_ of 0.4, indicating partial PerR derepression.

In addition, the H_2_O_2_ resistance of WT cells was 1.5-4-fold further enhanced during the stationary phase, whereas the Δ*katA* mutant did not survive 40 mM H_2_O_2_ during the log and stationary phases **(Fig. 1G)**. These data indicate that KatA contributes also to the stationary phase H_2_O_2_ resistance in *S. aureus*. In contrast, the peroxiredoxin-deficient *S. aureus* Δ*tpx* and Δ*bcp* mutants were not impaired in growth or survival after exposure 10 mM and 40 H_2_O_2_ **(Fig. 1C-E)**. However, the Δ*ahpC* mutant displayed an increased H_2_O_2_ resistance **(Fig. 1B, E)**, which is mediated by the PerR-dependent up-regulation of KatA in the Δ*ahpC* mutant as shown previously (Cosgrove et al., 2007). The derepression of the PerR regulon genes *katA* and *dps* in the Δ*ahpC* mutant was confirmed using Northern blots **(Fig. 1H)**. The growth phenotype of the *ahpC* mutant could be fully restored to WT level upon exposure to 10 mM H_2_O_2_ in the *ahpC* complemented strain **(Fig. S2A)**. However, in survival assays the resistance to 40 mM H_2_O_2_ could be only partially complemented, but significantly decreased at the 4h time point compared to the *ahpC* mutant **(Fig. S2D)**. This partial complementation of the H_2_O_2_ resistance in survival assays might be caused by the lower plasmid-borne AhpC expression compared to the highly abundant AhpC in WT cells as previously observed in the proteome (Wolf et al., 2008). To confirm whether KatA and AhpC play additive roles in H_2_O_2_ resistance, the growth and survival of the Δ*ahpC*Δ*katA* double mutant was analyzed. In agreement with previous data, the Δ*ahpC*Δ*katA* mutant showed a slower aerobic growth **(Fig. S3A)** (Cosgrove et al., 2007) and displayed 3-fold and 13-fold reduced survival rates after exposure to 1 mM H_2_O_2_ as compared to the Δ*katA* mutant **(Fig. 1F)**.

Taken together, these results indicate that KatA plays the major role to confer the strong H_2_O_2_ resistance to aerobic *S. aureus* cells during the log and the stationary phases, whereas AhpC contributes to a minor part to the H_2_O_2_ resistance during the aerobic growth. Thus, KatA was identified as major determinant of the H_2_O_2_ resistance in growing and non-growing *S. aureus* cells.

### The Δ*katA* mutant shows a strong oxidative shift in the *E*_BSH_ by H_2_O_2_ and is impaired to regenerate the reduced state as revealed by the Brx-roGFP2 biosensor

To monitor the changes in the *E*_BSH_ in the catalase and peroxiredoxin deficient mutants, we measured the Brx-roGFP2 biosensor responses after H_2_O_2_ stress in the WT and mutant strains. Consistent with previous data, Brx-roGFP2 was fast and strongly oxidized by 100 mM H_2_O_2_ in the *S*. _BSH_ *aureus* WT and cells could quickly recover the reduced state of *E*_BSH_ within two hours **(Fig**. _BSH_ **2A)**. However, due to its high resistance, no increased biosensor oxidation was measured in WT cells by 1 mM H_2_O_2_ **(Fig. 2B)**. In contrast to WT cells, the biosensor was fully and constitutively oxidized by 1 and 100 mM H_2_O_2_ in the Δ*katA* mutant, indicated by an impaired regeneration of reduced *E*_BSH_ **(Fig. 2A, B)**. The *katA* mutant was only able to recover the reduced state of *E*_BSH_ after treatment with 0.4 mM H_2_O_2_ **(Fig. 2C)**, suggesting that this low H_2_O_2_ level might be detoxified by AhpC. In support of this hypothesis, the Brx-roGFP2 biosensor was fast oxidized and strongly delayed in recovery of reduced *E*_BSH_ after exposure to 0.4 mM H_2_O_2_ in the Δ*ahpC*Δ*katA* double mutant **(Fig. S3B)**. In contrast, the Δ*ahpC*, Δ*tpx* and Δ*bcp* mutants showed similar H_2_O_2_ responses and regeneration of reduced *E*_BSH_ compared to the WT **(Fig. 2D-F)**. The biosensor results confirmed the hypersensitivities of the Δ*katA* and Δ*ahpC*Δ*katA* mutants towards H_2_O_2_ stress, supporting that KatA is responsible for the fast detoxification of 100 mM of H_2_O_2_ and regeneration of *E*_BSH_ in *S. aureus* WT cells, while AhpC can only detoxify low levels of 0.4 mM H_2_O_2_ in the absence of KatA. However, the peroxiredoxins Tpx and Bcp are not essential for H_2_O_2_ detoxification in *S. aureus* WT cells.

**Fig. 2.**
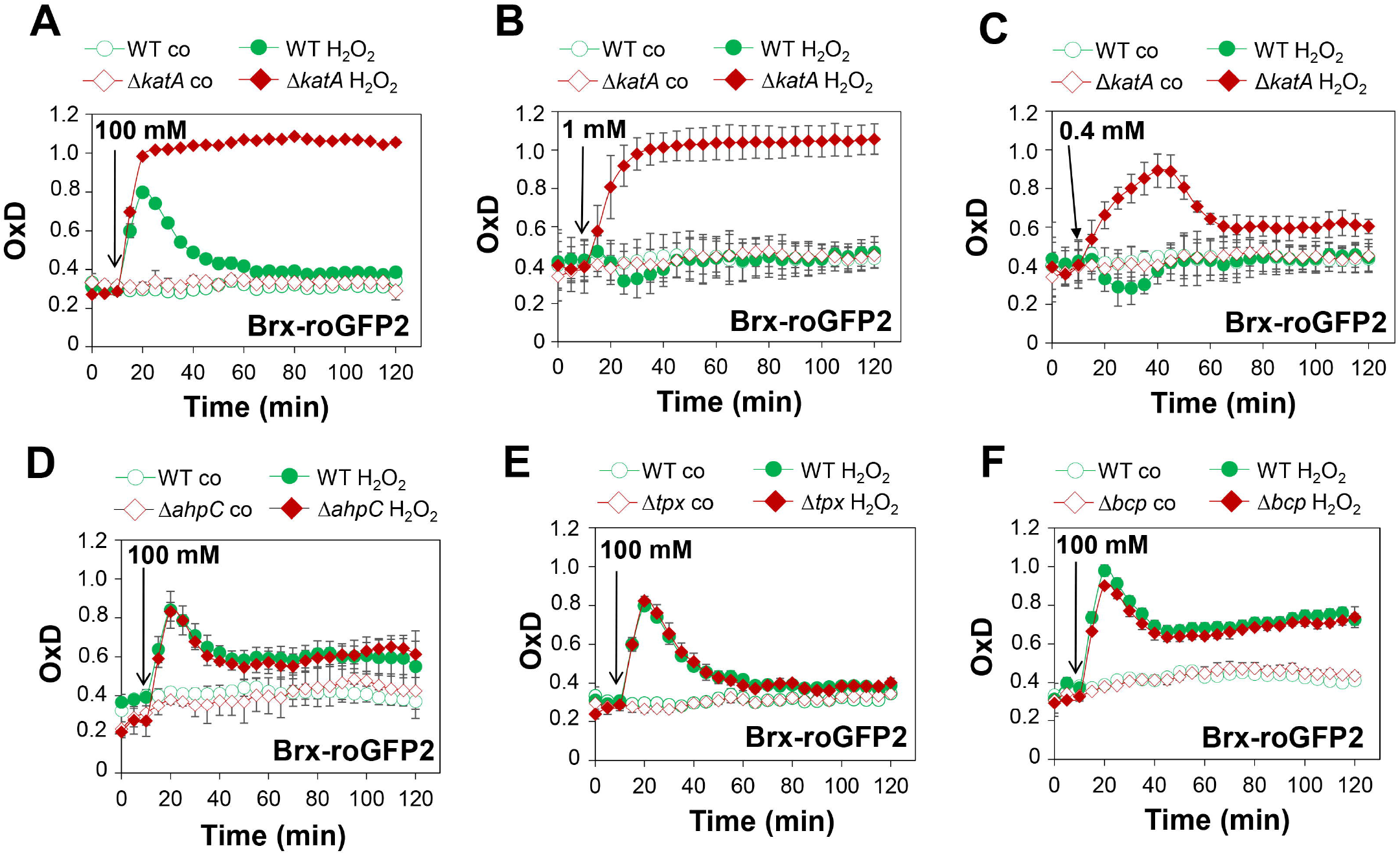
**The** Δ***katA* mutant can only recover from 0.4 mM H**_**2**_**O**_**2**_ **stress, but is unable to regenerate the reduced state of *E***_**BSH**_ **upon exposure to 1 mM H**_**2**_**O**_**2**_ **as revealed by Brx-roGFP2 measurements in *S. aureus*.** **(A-E)** Brx-roGFP2 biosensor responses to 100, 1 and 0.4 mM H_2_O_2_ were monitored in the *S. aureus* COL WT, Δ*katA* **(A, B, C)**, Δ*ahpC* **(D)**, Δ*tpx* **(E)** and Δ*bcp* mutants **(F)**. The oxidation degree (OxD) of the Brx-roGFP2 responses were calculated based on the 405/488 nm excitation ratios and normalized to fully reduced and oxidized controls. Mean values and SD of the OxD values are presented from 3 biological replicates.

### *S. aureus* shows a KatA-dependent microaerophilic H_2_O_2_ priming to acquire an improved resistance towards lethal H_2_O_2_ doses

The previous data revealed that KatA confers the strong H_2_O_2_ resistance during the aerobic growth. However, the role of KatA in priming of *S. aureus* for improved H_2_O_2_ resistance during microaerophilic conditions has not been investigated. Thus, the *S. aureus* WT and the Δ*katA* mutant were grown under microaerophilic conditions to the log phase and primed with 0.1 mM H_2_O_2_ for 30 min, followed by triggering with 10 mM H_2_O_2_ **(Fig. 3A)**. The growth and survival was analysed for naïve (C), primed (P), primed and triggered (PT) and triggered cells (T) **(Fig. 3A)**. The primed *S. aureus* WT and Δ*katA* mutant (P) were not impaired in growth and survival under 0.1 mM H_2_O_2_ **(Fig. 3B-E)**. However, the primed and triggered WT (PT) could acquire an improved resistance towards the triggering stimulus of 10 mM H_2_O_2_, compared to the triggering only state (T) **(Fig. 3B, D)**. Specifically, PT cells showed survival rates of 52-72%, whereas T cells were almost killed and survived only to <0.07 % after 10 mM H_2_O_2_ treatment. This indicates that *S. aureus* is primable for improved oxidative stress resistance during the microaerophilic growth **(Fig. 3B, D)**. However, in contrast to the WT, the primed Δ*katA* mutant strain was unable to acquire the improved resistance towards otherwise lethal doses of 10 mM H_2_O_2_ under microaerophilic conditions **(Fig. 3C, E)**. Both PT and T cells of the Δ*katA* mutant were strongly impaired in growth and completely killed after treatment with 10 mM H_2_O_2_ as triggering stimulus **(Fig. 3C, E)**. However, due to its lower catalase activity by plasmid-based constitutive KatA expression **(Fig. S1D)**, the *katA* complemented strain did not recovered the improved H_2_O_2_ resistance upon microaerophilic priming, resulting in growth inhibition and similar killing of PT and T cells **(Fig. S4)**. Overall, these results indicate that the catalase KatA is responsible for *S. aureus* priming for improved resistance towards upcoming lethal H_2_O_2_ stress under microaerophilic conditions. Taken together, KatA confers the constitutive resistance during aerobic growth and prepares *S. aureus* for up-coming future oxidative stress under microaerophilic conditions.

**Fig. 3.**
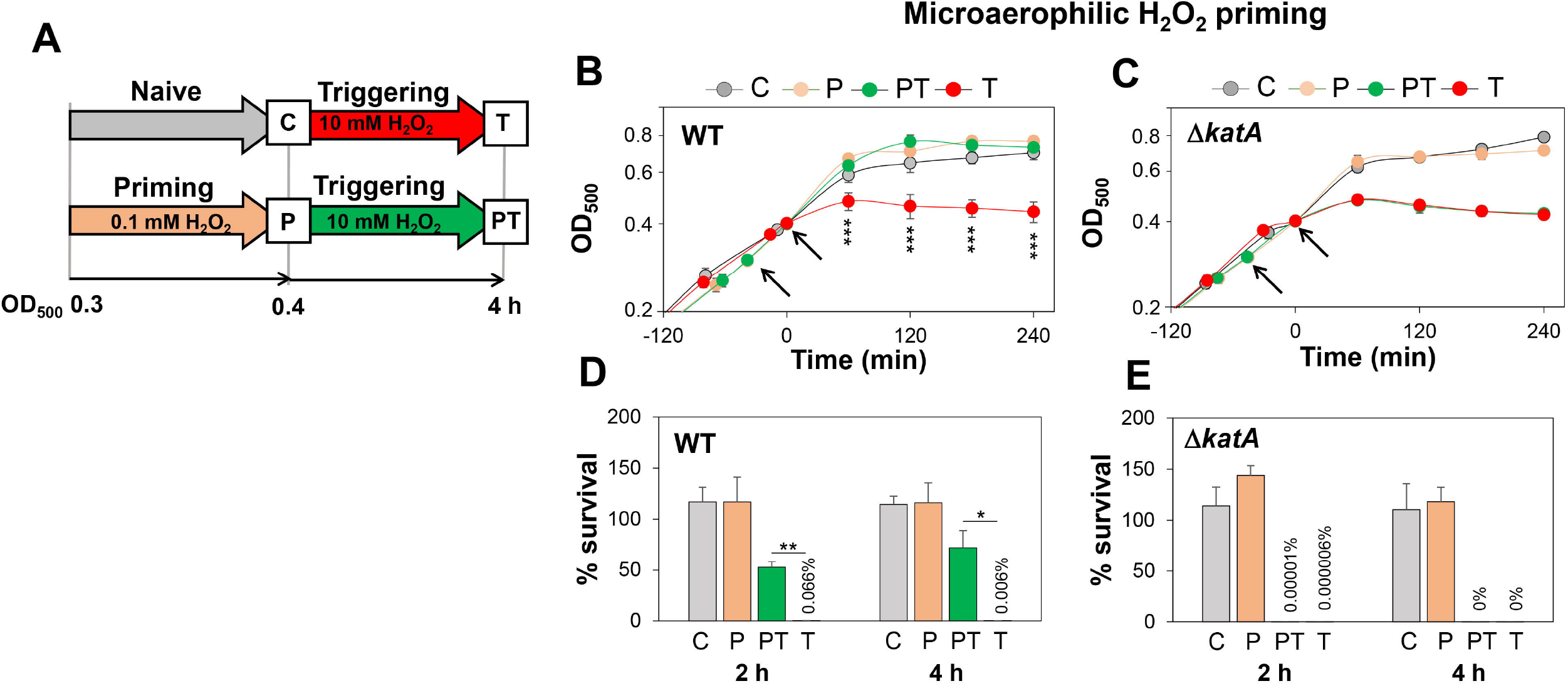
Microaerophilic H_2_O_2_ priming confers an improved resistance against otherwise lethal H_2_O_2_ doses in *S. aureus*, which depends on KatA. **(A)** Setup for microaerophilic priming and triggering experiments. The *S. aureus* WT and Δ*katA* mutant strains were grown microaerophilic and primed during the log phase with 0.1 mM H_2_O_2_ for ∼30 min (P), followed by treatment with 10 mM H_2_O_2_ as triggering stimulus (PT). The growth curves **(B, C)** and survival rates **(D, E)** were measured in naïve (C), primed (P), primed and triggered (PT) and triggered only cells (T). The survival rates were calculated after 2 and 4 h of H_2_O_2_ stress relative to untreated control cells. The results are from 3-4 biological replicates. Error bars represent the SD. The statistics was calculated using the Student’s unpaired two-tailed *t*-test by graph prism. Symbols are: *p < 0.05,**p ≤ 0.01 and ***p ≤ 0.001.

### *S. aureus* is not primable for improved H_2_O_2_ resistance during aerobic growth

Next, priming and triggering experiments were performed in aerobically grown *S. aureus* cells to analyse whether the constitutive H_2_O_2_ resistance can be further enhanced in primed cells **(Fig. 4)**. First, we used the same H_2_O_2_ doses for priming (0.1 mM) and triggering (10 mM) experiments, as applied in the microaerophilic experiments **(Fig. 4A)**. As expected, there was no difference in growth and survival between PT and T cells after exposure to 10 mM H_2_O_2_ during the aerobic growth **(Fig. 4B, C)**. Both PT and T cells were similar resistant and fully survived the 10 mM H_2_O_2_ triggering dose. Thus, the constitutive resistance of aerobic *S. aureus* cells towards 10 mM H_2_O_2_ could be not further enhanced by pre-exposure to the priming stimulus of 0.1 mM H_2_O_2_ **(Fig. 4B, C)**. As shown before, this constitutive H_2_O_2_ resistance of aerobically grown *S. aureus* cells was dependent on KatA **(Fig. 1A, E, F)**. Next, we increased the H_2_O_2_ doses for priming (1 mM H_2_O_2_) and triggering (40 mM) of *S. aureus* during the aerobic growth **(Fig 4D)**. However, also higher priming doses could not improve the growth and survival of PT cells in response to the subsequent 40 mM H_2_O_2_ stress compared to the T cells treated with 40 mM H_2_O_2_ only **(Fig. 4E, F)**. Both PT and T cells were strongly impaired in growth under 40 mM H_2_O_2_ stress and showed survival rates of <10% after 2 h. Small survival differences of 4% were observed after 4h in T versus PT cells, but these were not reproducible at the 2h time point **(Fig. 4F)**. In conclusion, these different priming setups support that *S. aureus* is not primable for further enhanced H_2_O_2_ resistance under aerobic conditions. We hypothesize that ROS production during aerobic respiration acts as a priming stimulus to induce expression of KatA, which confers the high H_2_O_2_ resistance.

**Fig. 4.**
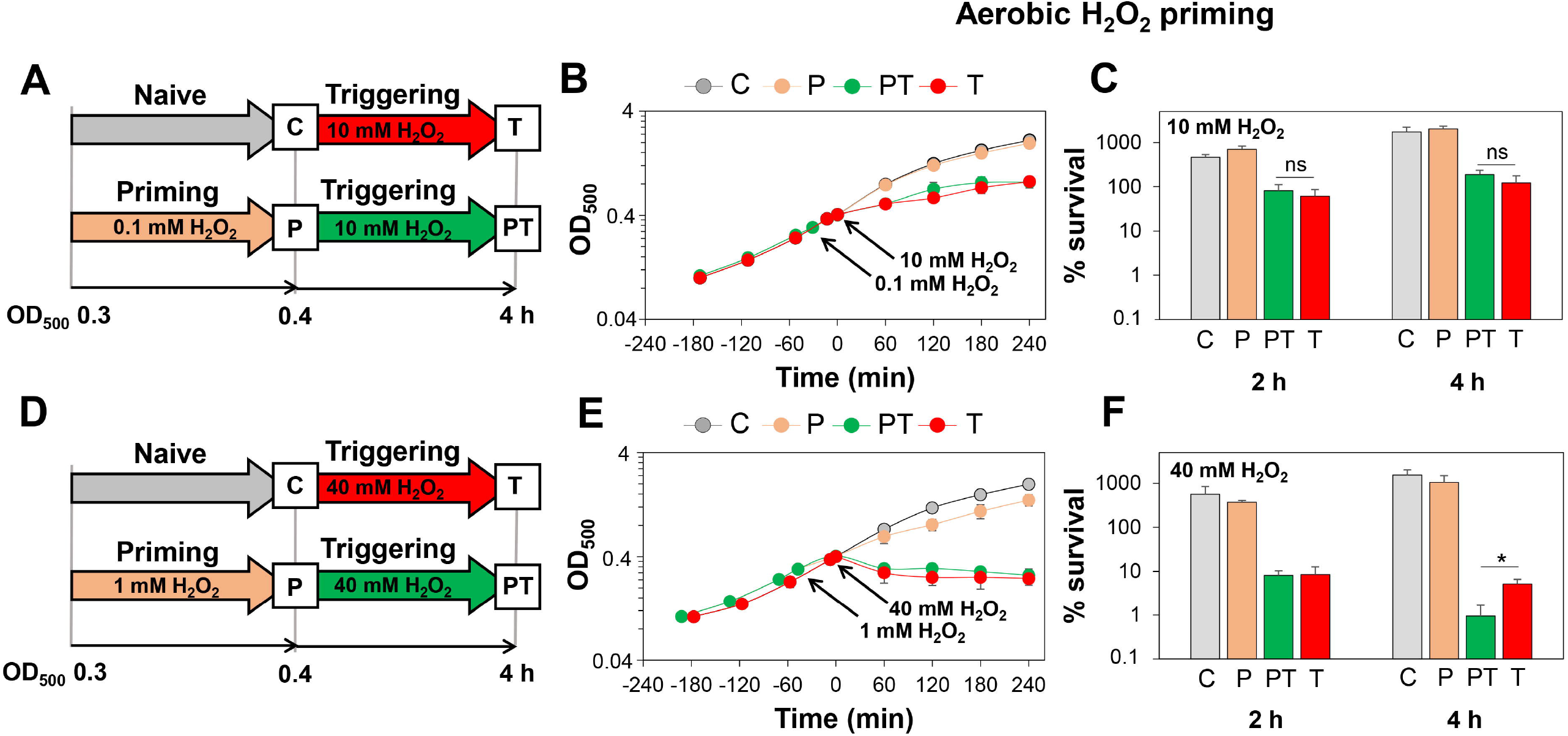
*S. aureus* is not primable for improved H_2_O_2_ resistance during aerobic growth. **(A, D)** Setup for aerobic priming and triggering experiments. The aerobically grown *S. aureus* WT was primed during the log phase with either 0.1 mM H_2_O_2_ **(A)** or 1 mM H_2_O_2_ **(D)** for ∼30 min (P) and subsequently treated with either 10 mM H_2_O_2_ **(A)** or 40 mM H_2_O_2_ **(D)**, respectively, as triggering stimuli (PT). The growth curves **(B, E)** and survival rates **(C, F)** were measured in both setups for naïve (C), primed (P), primed and triggered (PT) and triggered only cells (T). Survival rates of the H_2_O_2_ -treated cells were calculated relative to the untreated control. The results are from 3-4 biological replicates. Error bars represent the SD. The statistics was calculated using the Student’s unpaired two-tailed *t*-test by graph prism. Symbols are: ^ns^p > 0.05 and *p < 0.05.

### Microaerophilic H_2_O_2_ priming causes increased transcription of *katA* and elevated KatA activity, which confers improved resistance towards lethal H_2_O_2_ doses in *S. aureus*

Northern blot analyses were used to study whether microaerophilic H_2_O_2_ priming induces transcription of the PerR-dependent *katA* gene and the *ahpCF* operon in *S. aureus* **(Fig. 5)**. The results revealed that transcription of *katA* is significantly 1.8-fold up-regulated upon microaerophilic priming with 0.1 mM H_2_O_2_ **(Fig. 5B, D)**, whereas the basal level of *katA* transcription was already higher under aerobic conditions and could be not further induced during aerobic H_2_O_2_ priming **(Fig. 5C, E)**. Transcription of the *ahpCF* operon was not significantly induced during microaerophilic priming **(Fig. 5B, G)**. However, transcription of *katA* and *ahpCF* was strongly reduced after triggering by 10 mM H_2_O_2_ under microaerophilic conditions, since the triggering dose was found lethal **(Fig. 5B, D, G)**. Transcription of *katA* decreased even in the PT cells compared to P cells, which highlights the high efficiency of the KatA protein for fast removal of H_2_O_2_ in PT cells **(Fig. 5B, D)**. Under aerobic conditions, PT and T cells did not show significantly enhanced transcription of *katA* and *ahpCF*, supporting that the constitutive H_2_O_2_ resistance could not be further enhanced by pre-exposure to the priming dose **(Fig. 5C, E, H)**. The *katA* transcript levels could be confirmed by catalase activities during the microaerophilic and aerobic priming experiments **(Fig. 5F)**. Specifically, the basal activity of KatA was very low during the microaerophilic growth, whereas aerobic cells showed a constitutive high catalase activity. The KatA activity could be only enhanced upon microaerophilic priming, but not during aerobic priming due to the high constitutive resistance **(Fig. 5F)**. Further consistent with the Northern blots, the KatA activity decreased strongly in PT and T cells during microaerophilic priming. Together, these results revealed that microaerophilic priming induces *katA* transcription and KatA activity, which confers improved resistance towards otherwise lethal H_2_O_2_ stress.

**Fig. 5.**
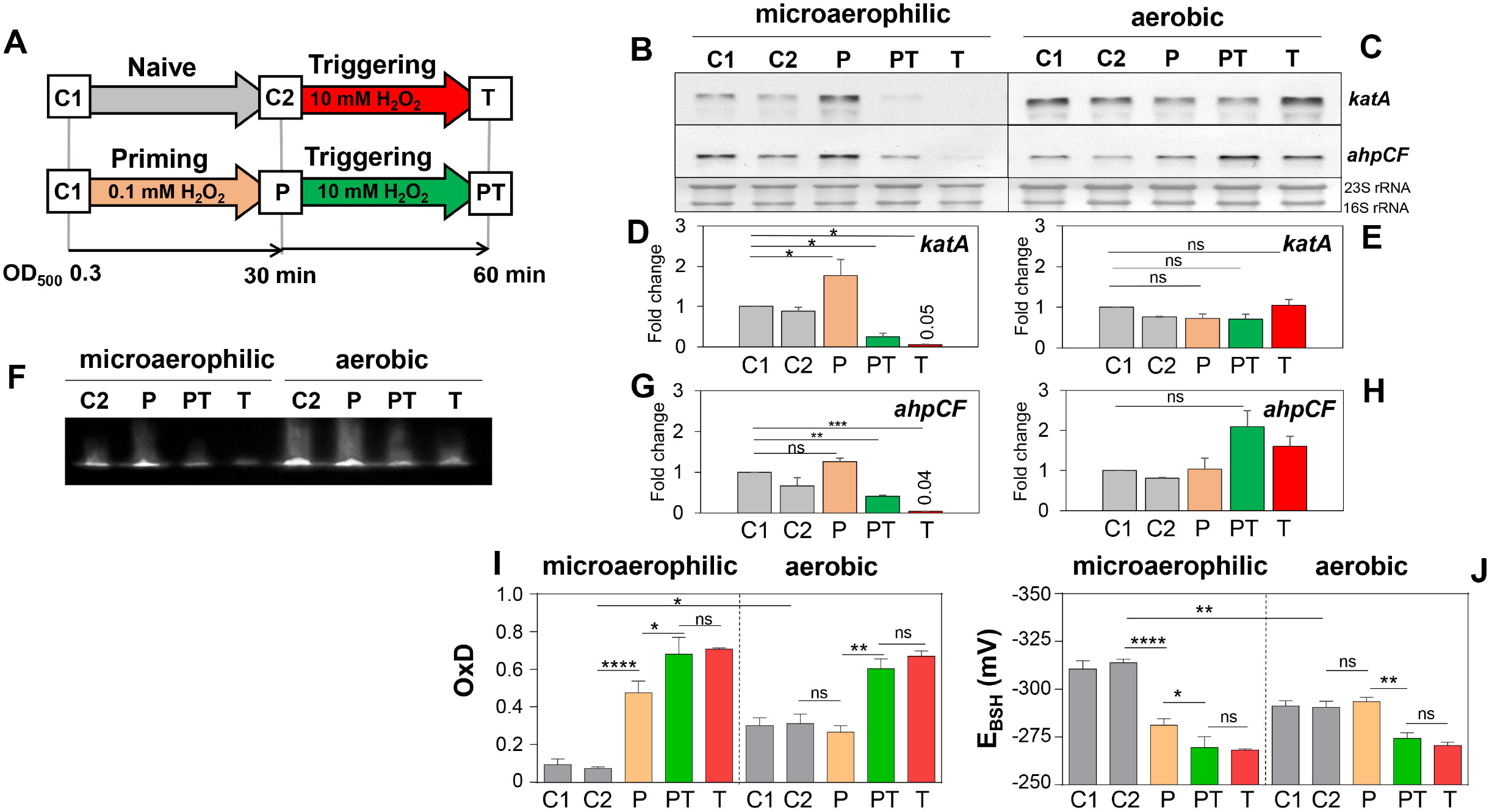
**Microaerophilic H**_**2**_**O**_**2**_ **priming leads to increased *katA* transcription and KatA activity, which mediates the improved H**_**2**_**O**_**2**_ **resistance in *S. aureus*. In addition, the Brx-roGFP2 biosensor is fast oxidized upon microaerophilic H**_**2**_**O**_**2**_ **priming, but not under aerobic conditions.** **(A)** The setup for microaerophilic and aerobic priming and triggering experiments includes priming with 0.1 mM H_2_O_2_ and triggering with 10 mM H_2_O_2_. **(B, C)** To analyze transcription of *katA* and *ahpCF* using Northern blots, RNA was isolated from *S. aureus* WT cells in the naïve (C1, C2), primed (P), primed and triggered (PT) and triggered (T) states. The band intensities of the *katA* **(D, E)** and *ahpCF* specific transcripts **(G, H)** were quantified from 2 biological replicates using ImageJ and error bars represent the SD. The statistics was calculated using the ordinary one-way ANOVA and Dunnet’s multiple comparisons test using graph prism. Symbols are defined as: ns p>0.05; *p≤0.05; **p≤0.01 and ***p≤ 0.001. **(F)** The catalase activity was analyzed in cell extracts of the *S. aureus* WT during microaerophilic and aerobic H_2_O_2_ priming at C1, C2, P, PT and T using native PAGE and diaminobenzidine staining. **(I, J)** The response of the Brx-roGFP2 biosensor was measured in *S. aureus* COL grown in LB medium under microaerophilic and aerobic conditions in naïve (C1, C2), primed (P), primed and triggered (PT) and triggered (T) cells. The C1, C2 samples were harvested at OD_500_ of 0.3 and 0.4, respectively, and the P, PT and T cells were harvested after 10 min of H_2_O_2_ exposure (maximum of biosensor oxidation). Samples were blocked 10 mM NEM and the fluorescence excitation maxima were measured at 405 and 488 nm using the microplate reader. OxD values and *E*_BSH_ of Brx-roGFP2 were calculated using the 405/408 nm excitation ratio as described previously. Mean values of 3 biological replicates are shown, error bars represent the SD and p-values were calculated using a Student’s unpaired two-tailed t-test by the graph prism software. Symbols are defined as: ns p>0.05; *p≤0.05 and **p≤0.01.

### Microaerophilic priming leads to a strong oxidative shift in the *E*_BSH_

We were interested in the *E*_BSH_ differences between aerobic and microaerophilic growth conditions, supporting the enhanced ROS levels in aerobic cells as endogenous priming stimulus. Moreover, we aimed to analyse the Brx-roGFP2 biosensor response upon microaerophilic priming and triggering to investigate if the KatA induction upon microaerophilic priming is accompanied by a change in the Brx-roGFP2 biosensor oxidation. The comparison of the basal biosensor oxidation revealed a more reducing basal OxD of 0.1 and an *E*_BSH_ of -310 mV during microaerophilic growth (C1) compared to the OxD of 0.3 and the *E*_BSH_ of -291 mV during aerobic growth (C1) **(Fig. 5I, J)**. Thus, the higher basal oxidation in aerobic cells accounts for the increased ROS level due to aerobic respiration.

Upon microaerophilic priming, the Brx-roGFP2 biosensor showed a 5-fold increased OxD and an oxidized *E*_BSH_ of -281 mV, which was further oxidized in PT and T cells. In contrast, the biosensor did not respond to aerobic priming and showed an increased oxidation only in aerobic PT and T cells **(Fig. 5I, J)**. These results support that microaerophilic priming leads to an oxidative shift of the *E*_BSH_ from -310 mV to -281 mV, resulting in increased catalase expression to prime the cells for improved H_2_O_2_ resistance. Due to the higher basal oxidation in aerobic cells, priming did not change the high *E*_BSH_ of - 291 mV, which is consistent with the constitutive KatA expression **(Fig. 5I, J)**. These results on the *E*_BSH_ differences of ∼20 mV between microaerophilic and aerobic cells strongly support that respiratory ROS primes aerobic cells for the constitutive H_2_O_2_ resistance.

### The peroxiredoxins AhpC, Tpx and Bcp mediate CHP resistance in *S. aureus*

While the role of KatA in aerobic H_2_O_2_ resistance and microaerophilic H_2_O_2_ priming has been clearly revealed, the peroxiredoxins AhpC, Tpx and Bcp could function also in the resistance to organic hydroperoxides in *S. aureus*. Using growth and survival assays, the phenotypes of the catalase and peroxiredoxin-deficient mutants were analyzed after CHP treatment **(Fig. 6)**. While the growth of the Δ*katA* and Δ*bcp* mutants was not affected by 0.15 mM CHP stress, the CHP-treated Δ*ahpC* and Δ*tpx* mutant showed slightly reduced growth rates **(Fig. 6A-D)**. However, the Δ*ahpC*, Δ*tpx* and Δ*bcp* mutants showed significantly decreased survival rates under CHP stress at the 4 h time point, supporting that the peroxiredoxins confer protection against CHP stress in *S. aureus* **(Fig. 6E)**. In addition, the Δ*katA* mutant showed a slightly higher CHP resistance, which could be not reverted to WT levels in the *katA* complemented strain, probably due to the partial complementation by plasmid-based KatA expression **(Fig. 6E)**. However, the CHP-sensitive survival phenotypes of the peroxiredoxin-deficient mutants could be restored to WT levels in the *ahpC, tpx* and *bcp* complemented strains **(Fig. S2E, F)**.

**Fig. 6.**
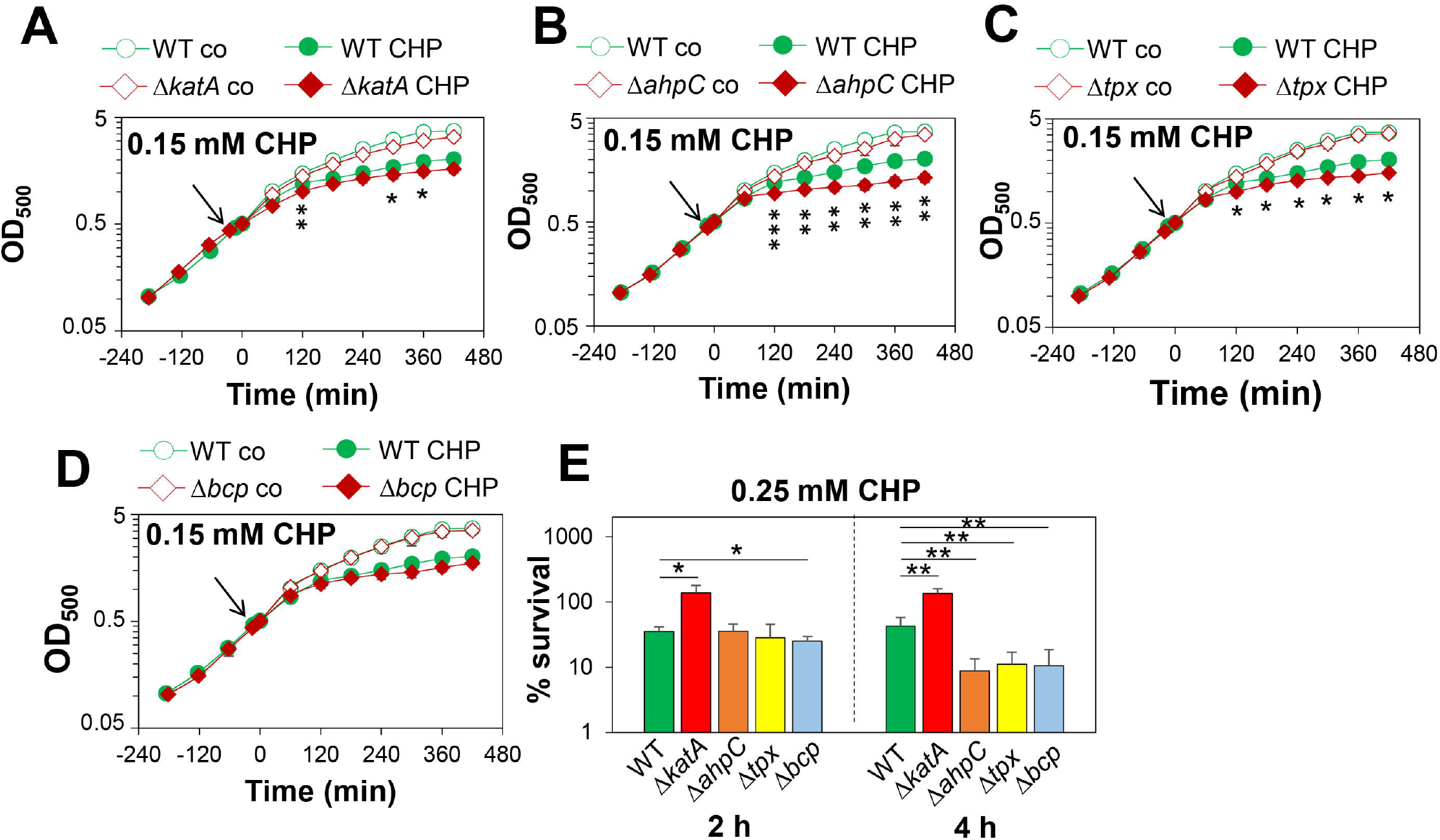
The peroxiredoxins AhpC, Tpx and Bcp contribute to CHP resistance in *S. aureus*. **(A-D)** Growth phenotypes were analyzed of the *S. aureus* COL WT, Δ*katA* **(A)**, Δ*ahpC* **(B)**, Δ*tpx* **(C)** and Δ*bcp* mutants **(D)** in RPMI medium before (co) after exposure to 0.15 mM CHP stress at an OD_500_ of 0.5. **(E)** The survival rates of the *S. aureus* strains were calculated 2 and 4 h after exposure to 0.25 mM CHP relative to the untreated control, which was set to 100%. Mean values and SD of 4-6 biological replicates are presented. The statistics was calculated using the Student’s unpaired two-tailed t-test by graph prism. Symbols are: ^ns^p > 0.05, *p ≤ 0.05, **p ≤ 0.01 and ***p ≤ 0.001.

### The peroxiredoxin-deficient Δ*ahpC*, Δ*tpx* and Δ*bcp* mutants are delayed in CHP detoxification as revealed by Brx-roGFP2 measurements

To investigate the impact of the peroxiredoxins in the maintenance of the reduced state of *E*_BSH_ in *S. aureus*, Brx-roGFP2 biosensor measurements were performed after 0.5 mM CHP stress **(Fig 7)**. The Brx-roGFP2 biosensor was similarly fast oxidized by 0.5 mM CHP in the WT, Δ*katA*, Δ*ahpC*, Δ*tpx* and Δ*bcp* mutants. However, while the WT and Δ*katA* mutant could regenerate the reduced state of *E*_BSH_ within 2 hours, the Δ*ahpC* mutant was unable to regenerate the basal level of *E*_BSH_ during the recovery phase from CHP stress **(Fig. 7A,B)**. In addition, both Δ*bcp* and Δ*tpx* mutants showed a significant delay in recovery of the reduced state of *E*_BSH_ upon CHP stress as compared to the WT **(Fig. 7C,D)**. These biosensor results support that the peroxiredoxins AhpC, Tpx and Bcp are important for CHP detoxification and contribute to regeneration of *E*_BSH_ during the recovery phase from CHP stress in *S. aureus*. In contrast, KatA does not contribute to CHP detoxification and resistance in *S. aureus*.

**Fig. 7.**
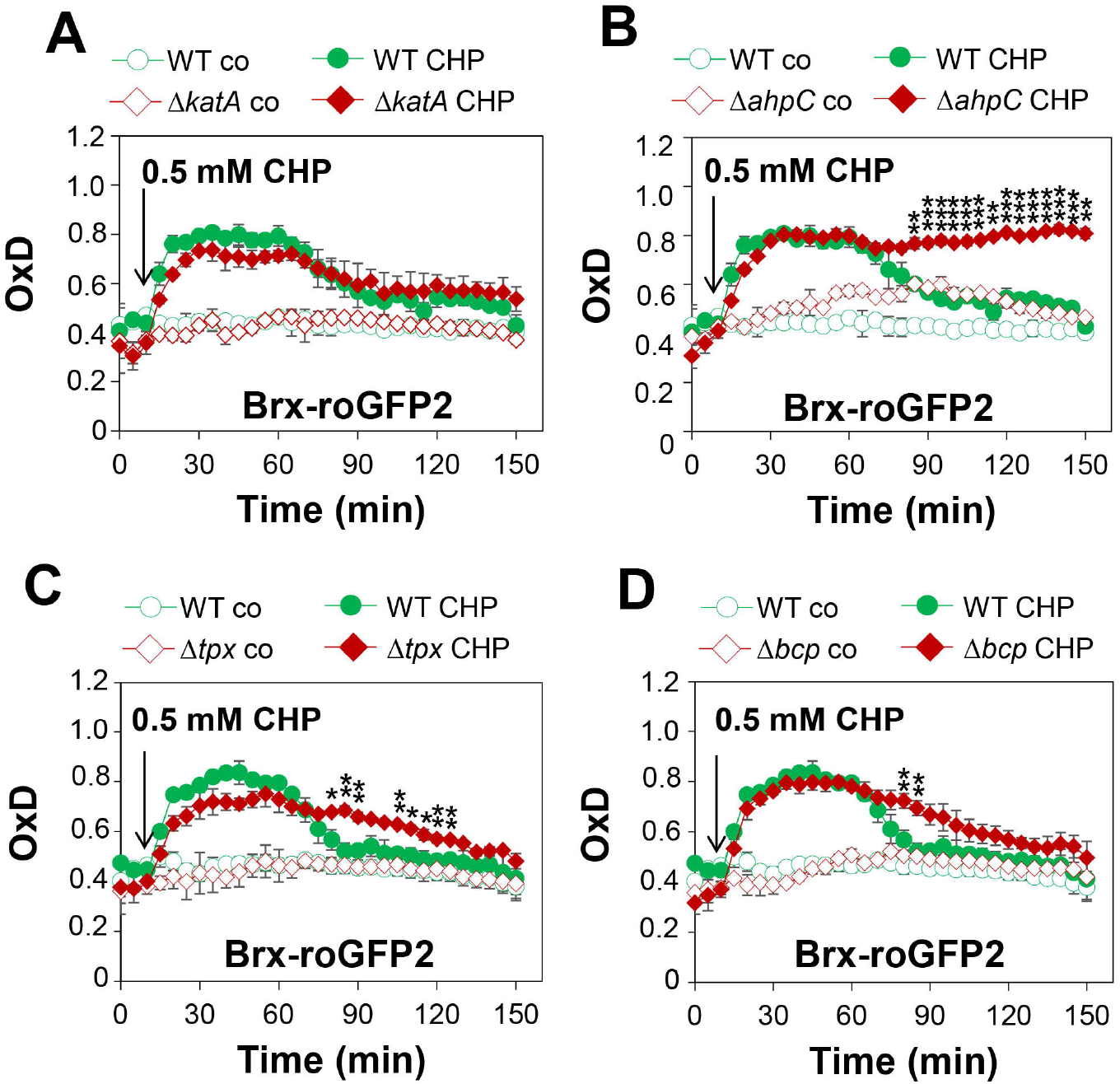
AhpC, Tpx and Bcp function in CHP detoxification to regenerate the reduced *E*_BSH_ during the recovery phase as revealed by the Brx-roGFP2 biosensor. **(A-D)** Monitoring Brx-roGFP2 responses to 0.5 mM CHP in *S. aureus* COL WT, Δ*katA* **(A)**, Δ*ahpC* **(B)**, Δ*tpx* **(C)** and Δ*bcp* mutants **(D)**. The oxidation degree (OxD) of the Brx-roGFP2 response was calculated based on the 405/488 nm excitation ratio and normalized to fully reduced and oxidized controls. Mean values and SD of 3 biological replicates are shown. The statistics was calculated using the Student’s unpaired two-tailed t-test by graph prism. Symbols are: ^ns^p > 0.05, *p ≤ 0.05, **p ≤ 0.01 and ***p ≤ 0.001.

### Catalases and peroxiredoxins do not contribute to protection against HOCl stress

Previous transcriptome analyses revealed an increased transcription of the PerR regulon under HOCl stress in *S. aureus* (Loi et al., 2018). Thus, growth curves and survival assays were used to analyse the phenotypes of the Δ*katA*, Δ*ahpC*, Δ*tpx* and Δ*bcp* mutants under HOCl stress. However, none of these mutants showed significant defects in growth or survival after HOCl stress compared to the WT, indicating that the catalase and peroxiredoxins do not contribute to HOCl detoxification and resistance in *S. aureus* **(Fig. S5A-E)**.

## DISCUSSION

In this manuscript, we have investigated the roles of the catalase and peroxiredoxins in the peroxide stress resistance and priming of *S. aureus* during aerobic and microaerophilic growth. Our results revealed that *S. aureus* is not primable towards improved H_2_O_2_ resistance during the aerobic growth due to its constitutive H_2_O_2_ resistance, which was shown to be dependent on the catalase KatA. Moreover, we found that microaerophilic *S. aureus* cells are primable to acquire an enhanced resistance towards lethal H_2_O_2_ doses, which was also mediated by KatA. In addition, the peroxiredoxins AhpC, Tpx and Bcp were shown to contribute to the CHP detoxification and resistance in *S. aureus* to ensure the maintenance of the redox balance upon recovery from stress.

The roles of KatA and AhpC in the peroxide resistance of *S. aureus* have been previously demonstrated (Cosgrove et al., 2007). In this work, we additionally used Brx-roGFP2 biosensor measurements, showing the impact of KatA and AhpC on the level of H_2_O_2_ detoxification and regeneration of reduced *E*_BSH_ under oxidative stress. Without KatA, *S. aureus* cells are highly sensitive to oxidants and only able to remove low doses of 0.4 mM H_2_O_2_, which were shown to be detoxified by AhpC. In addition, the Δ*ahpC*Δ*katA* double mutant showed an increased sensitivity towards H_2_O_2_ stress in survival assays as compared to the Δ*katA* mutant. These data confirm that KatA and AhpCF have compensatory roles in H_2_O_2_ resistance to ensure the survival of *S. aureus* under oxidative stress (Cosgrove et al., 2007). Expression of *katA, bcp* and the *ahpCF* operon is controlled by the peroxide-sensing PerR repressor, which is inactivated by H_2_O_2_ due to Fe^2+^-catalyzed histidine oxidation in *S. aureus* (Horsburgh et al., 2001;Lee and Helmann, 2006;Ji et al., 2015). Due to PerR derepression in the Δ*ahpC* mutant, *katA* and *dps e*xpression were elevated in the Δ*ahpC* mutant as confirmed here using Northern blots and shown previously (Cosgrove et al., 2007). Thus, the higher KatA expression level in the Δ*ahpC* mutant mediates the enhanced H_2_O_2_ resistance, confirming previous results in *S. aureus* and *B. subtilis* (Antelmann et al., 1996;Bsat et al., 1996;Cosgrove et al., 2007).

In addition, we showed that aerobically grown *S. aureus* acquire an improved resistance to H_2_O_2_ during the stationary phase, which also depends on KatA. The oxidative stress resistance was also enhanced during the stationary phase in other bacteria, such as *B. subtilis* and *E. coli* (Dowds et al., 1987;Farr and Kogoma, 1991). In addition, KatA activity was elevated during the stationary phase in *S. aureus* and *B. subtilis* (Loewen and Switala, 1987;Cosgrove et al., 2007). Altogether, our results on aerobic *S. aureus* cells revealed that KatA is the major player, which confers the strong constitutive H_2_O_2_ resistance during the log and stationary phases, while the peroxiredoxin AhpC plays an additional role to scavenge 0.4 mM H_2_O_2_ in the absence of KatA.

In this study, we further showed that *S. aureus* is primable with sub-lethal doses of 0.1 mM H_2_O_2_ to acquire an improved resistance towards otherwise lethal doses of 10 mM H_2_O_2_ during the microaerophilic growth. This microaerophilic H_2_O_2_ priming was found to be dependent on KatA, which was transcriptionally induced and showed a higher catalase activity upon challenge with the priming dose. These results are consistent with previous data showing increased KatA activity by exposure to 100 µM H_2_O_2_ during oxygen limitation (Ji et al., 2015). We showed that microaerophilic priming by KatA prepares the cells to survive better the lethal triggering stress, resulting in an improved growth and survival of *S. aureus*. In contrast, due to their constitutive H_2_O_2_ resistance, aerobic *S. aureus* cells were not primable to acquire higher resistance towards lethal H_2_O_2_ doses. Aerobic priming was not possible with 0.1 mM or 1 mM H_2_O_2_ since *katA* transcription and catalase activity were already elevated in naïve cells and could not be further increased in primed cells. The higher basal level of *E*_BSH_ of -291 mV in aerobic cells compared to the more reducing basal *E*_BSH_ of - 310 mV in microaerophilic cells strongly indicates the increased ROS level in aerobic cells due to aerobic respiration. These biosensor data support that *S. aureus* is primed by endogenous ROS generated by aerobic respiration to achieve their constitutive H_2_O_2_ resistance phenotype **(Fig. 8)**. Thus, the PerR regulon is already up-regulated during the aerobic growth due to ROS generated during aerobic respiration (Ji et al., 2015), resulting in the H_2_O_2_ resistant phenotype.

**Fig. 8.**
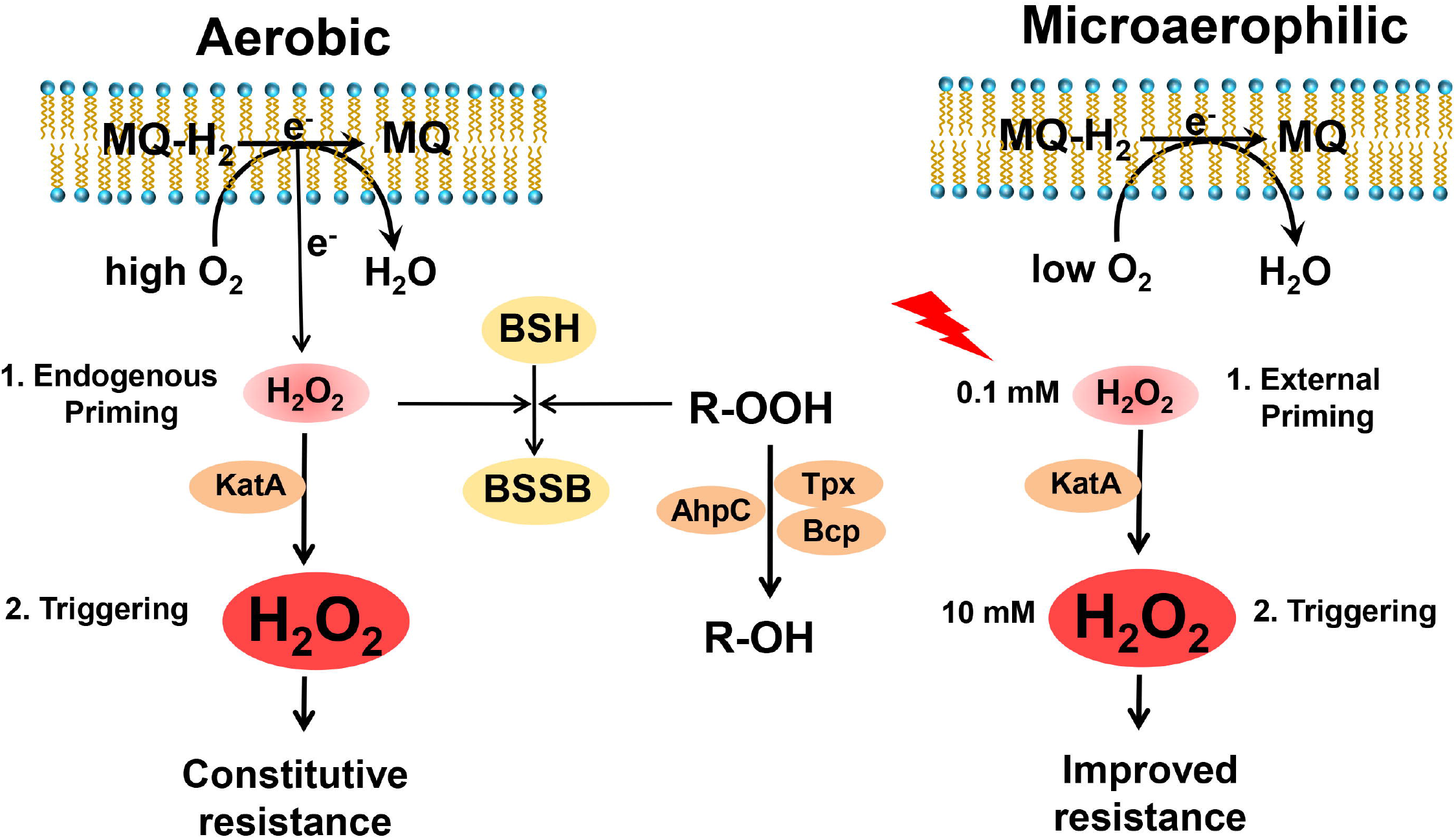
Summary schematics of microaerophilic H_2_O_2_ priming and constitutive aerobic H_2_O_2_ resistance in *S. aureus*. In this work, we showed that microaerophilic priming with 100 µM H_2_O_2_ induces improved resistance towards otherwise lethal doses of 10 mM H_2_O_2_ in *S. aureus*, while aerobic cells are already primed by ROS originating from aerobic respiration resulting in the constitutive high H_2_O_2_ resistance. The higher ROS levels during the aerobic growth were revealed by the ∼20 mV increased basal *E*_BSH_ compared to microaerophilic cells. The catalase KatA was shown to contribute to the microaerophilic H_2_O_2_ priming and aerobic H_2_O_2_ resistance in *S. aureus*, while the peroxiredoxins AhpC, Tpx and Bcp confer resistance towards organic hydroperoxides. In addition, KatA and the peroxiredoxins contribute to the regeneration of reduced *E*_BSH_ upon recovery from oxidative stress. MQ-H_2_ and MQ indicate reduced and oxidized menaquinone. BSH and BSSB are reduced bacillithiol and oxidized bacillithiol disulfide, respectively.

This up-regulation of the PerR regulon in *S. aureus* during the aerobic growth was caused by the hypersensitive PerR repressor, which is poised by very low endogenous levels of H_2_O_2_, generated during aerobic respiration (Ji et al., 2015). The endogenous H_2_O_2_ concentration in aerobic *E. coli* cells was determined as ∼50 nM (Seaver and Imlay, 2001). While PerR of *S. aureus* is hypersensitive to sense endogenous H_2_O_2_ levels during aerobic growth, PerR of *B. subtilis* is less sensitive and cannot sense such low H_2_O_2_ levels originating from aerobic respiration (Ji et al., 2015). Thus, *B. subtilis* can be primed during aerobic growth with 0.1 mM H_2_O_2_, leading to PerR inactivation and derepression of the PerR-controlled KatA, which confers an adaptive resistance against the otherwise lethal triggering dose of 10 mM H_2_O_2_ (Murphy et al., 1987;Antelmann et al., 1996;Engelmann and Hecker, 1996;Ji et al., 2015;Pinochet-Barros and Helmann, 2018). Similarly, aerobic priming is possible in *E. coli* and *S*. Typhimurium after exposure to 100 µM H_2_O_2_, leading to activation of the OxyR regulon, including the major catalase, which confers an adaptive and improved resistance towards lethal concentrations of 10 mM H_2_O_2_ (Christman et al., 1985;Farr and Kogoma, 1991;Åslund et al., 1999). In conclusion, priming of bacteria towards improved H_2_O_2_ resistance depends on the sensitivity of the redox-sensing peroxide regulators. While many bacteria harbor less sensitive H_2_O_2_ sensors and are primable by sub-lethal H_2_O_2_ doses during the aerobic growth, *S. aureus* PerR is hypersensitive to endogenous H_2_O_2_ levels during the aerobic growth (Ji et al., 2015), resulting in the constitutive aerobic H_2_O_2_ resistance and microaerophilic priming to acquire improved H_2_O_2_ resistance only in oxygen-limited conditions **(Fig. 8)**.

While KatA is responsible for detoxification of up-to 100 mM H_2_O_2_ and confers the high resistance to 100 mM H_2_O_2_ in aerobic *S. aureus* cells, the peroxiredoxin AhpC was shown to enable the detoxification of low levels of 0.4 mM H_2_O_2_, which could originate from the aerobic respiration. Thus, our results support that catalases are scavenger of high mM levels of H_2_O_2_, whereas peroxiredoxins can only detoxify physiological µM H_2_O_2_ levels. In addition, the peroxiredoxins AhpC, Tpx and Bcp were shown to confer protection against CHP stress in *S. aureus* cells. However, KatA and the peroxiredoxins are not involved in the defense against HOCl stress in *S. aureus*. Using Brx-roGFP2 biosensor measurements, the peroxiredoxin-deficient Δ*ahpC*, Δ*tpx* and Δ*bcp* mutants showed delayed regeneration of the reduced state of *E*_BSH_ after recovery from CHP stress. Thus, AhpC, Tpx and Bcp function in CHP detoxification and contribute to the maintenance of the cellular redox balance.The role of AhpC in OHP resistance has been previously demonstrated in *S. aureus* (Cosgrove et al., 2007). Similarly, the AhpC homologs of *B. subtilis, E. coli* and *S*. Typhimurium conferred resistance towards CHP stress (Storz et al., 1989;Antelmann et al., 1996). In *E. coli*, the Δ*tpx* and Δ*bcp* mutants were more sensitive towards OHP stress, indicating that these peroxiredoxins are more specific for reduction of organic peroxide substrates (Jeong et al., 2000;Cha et al., 2004). Kinetic assays of the *E* .*coli* Tpx protein demonstrated the substrate specificity towards alkyl hydroperoxides over H_2_O_2_ (Baker and Poole, 2003). Similarly, Bcp of *E. coli* has an 5-fold higher V_max_/K_m_ value for linoleic acid hydroperoxide as substrate compared to H_2_O_2_ (Jeong et al., 2000).

Taken together, we have shown that the catalase KatA is the major player of the aerobic H_2_O_2_ resistance in *S. aureus* and mediates priming to endogenous ROS level generated during aerobic respiration to confer the constitutive H_2_O_2_ resistance to the triggering stimulus in aerobic cells. In addition, KatA mediates microaerophilic priming by low H_2_O_2_ levels to prepare *S. aureus* cells for an improved and adaptive resistance against otherwise lethal H_2_O_2_ doses. In addition, the peroxiredoxins AhpC, Tpx and Bcp were shown to contribute to CHP detoxification and resistance to ensure survival and regeneration of the reduced *E*_BSH_ in *S. aureus* **(Fig. 8)**. In future studies, we aim to elucidate the functions of KatA and the peroxiredoxins in signal transduction, as redox-active chaperones and in cellular metabolism of *S. aureus*. Oxidative stress causes increased protein thiol-oxidation of many metabolic enzymes, leading to *S*-bacillithiolation and inactivation of the glycolytic glyceraldehyde 3-phosphate dehydrogenase GapDH in *S. aureus* (Deng et al., 2013;Imber et al., 2018). On the metabolome level, GapDH inactivation in *S. aureus* and yeast cells resulted in the redirection of the glycolytic flux to the pentose phosphate pathway, to ensure NADPH production as electron source for thiol-disulfide reductases to regenerate oxidized proteins (Ralser et al., 2007;Deng et al., 2013;Deng et al., 2014;Imber et al., 2018). In addition, H_2_O_2_ can damage iron-sulfur clusters of hydratases, such as aconitase, fumarate reductase and isopropylmalate isomerase, affecting the flux in the tricarboxylic acid cycle, aerobic respiration and branched chain amino acid synthesis (Varghese et al., 2003;Imlay, 2006;Jang and Imlay, 2007). Thus, it will be interesting to study the impact of KatA and the peroxiredoxins on the oxidative inactivation of redox-sensitive enzymes and the resulting changes in the cellular metabolome in *S. aureus* during the aerobic growth and under oxidative stress.

## ACKNOWLEDGMENTS

We are very grateful to Franziska Kiele for excellent technical assistance. We further would like to thank Quach Ngoc Tung and Lukas Olff for their support in mutant constructions and phenotype analyses. This work was supported by an ERC Consolidator grant (GA 615585) MYCOTHIOLOME and grants from the Deutsche Forschungsgemeinschaft (AN746/4-1 and AN746/4-2) within the SPP1710, by the SFB973 project C08 and the SFB/TR84 project B06 to H.A. We further acknowledge support by the Open Access Publication Initiative of the Freie Universität Berlin.

## AUTHOR CONTRIBUTIONS

N.L. and V.V.L. performed the experiments and analyzed the data of the manuscript. V.V.L. constructed the mutants and provided advise for biosensor measurements. N.L. wrote the initial draft of the manuscript. H.A. designed and supervised the experiments, provided funding and wrote the final manuscript. All authors edited and approved the final manuscript.

## Supplementary Figure legends

**Fig. S1. Complementation of KatA increases the H_2_O_2_ resistance in *S. aureus* (A-C), but the catalase activity in the *katA* complemented strain is lower compared to the WT (D)**. Growth curves of the *S. aureus* COL WT **(A)**, Δ*katA* mutant **(B)** and the *katA* complemented strain **(C)** after exposure to 0.4 and 1 mM H O stress at an OD_500_ of 0.5. Mean values and SD of four biological replicates are presented. **(D)** Catalase staining of aerobic *S. aureus* COL WT, the Δ*katA* mutant and the *katA* complemented strain after exposure to 10 mM H_2_O_2_ 500 at an OD of 0.5. Protein extracts were separated by native PAGE and the catalase activity was determined using diamidobenzidine staining as described in the Methods.

**Fig. S2. The CHP-sensitive phenotypes of the** Δ***ahpC***, Δ***tpx* and** Δ***bcp* mutants are abolished in the complemented strains. (A-C)** Growth curves of *S. aureus* COL WT, Δ*ahpC* and Δ*tpx* mutants and the complemented strains in RPMI medium before (co) after exposure to 10 mM H_2_O_2_ **(A)** and 0.15 mM CHP **(B, C). (D-F)** The survival rates were determined for the *S. aureus* COL WT, Δ*ahpC*, Δ*katA* and Δ*tpx* mutants and complemented strains after exposure to 40 mM H_2_O_2_ **(D)** and 0.25 mM CHP **(E, F)** based on CFU counts related to the untreated control, which was set to 100%. Mean values and SD of 4 biological replicates are presented. The statistics was determined using the Student’s unpaired two-tailed t-test by graph prism. Symbols are: ^ns^p > 0.05, *p ≤ 0.05, **p ≤ 0.01 and ***p ≤ 0.001.

**Fig. S3. The** Δ***katA***Δ***ahpC* double mutant is hypersensitive towards H_2_O_2_ and delayed in recovery of the reduced *E***_**BSH**_ **after 0.4 mM H_2_O_2_ (A)** Growth curves of *S. aureus* COL WT and the Δ*ahpC*Δ*katA* double mutant in RPMI medium before (co) and after exposure to 10 mM H_2_O_2_ at an OD_500_ of 0.5. The survival rates were calculated relative to the untreated control, which was set to 100%. Mean values and SD of 3-4 biological replicates are shown. The statistics was calculated using the Student’s unpaired two-tailed t-test by the graph prism software. Symbols are *p ≤ 0.05, **p ≤ 0.01 and ***p ≤ 0.001. **(B)** The Brx-roGFP2 biosensor in the *S. aureus* COL Δ*ahpC*Δ*katA* double mutant showed a fast oxidation and a delay in recovery of the reduced *E*_BSH_ after 0.4 mM H_2_O_2_. The oxidation degree (OxD) of the Brx-roGFP2 response was calculated based on the 405/488 nm excitation ratios and normalized to fully reduced and oxidized controls. Mean values and SD of the OxD values are presented from 3 biological replicates.

**Fig. S4. The *katA* complemented strain is not primable towards increased H_2_O_2_ resistance during microaerophilic growth. (A)** Setup for microaerophilic priming and triggering experiments. The *S. aureus katA* complemented strain was primed during the log phase with 0.1 mM H_2_O_2_ for ∼30 min (P) and subsequently treated with 10 mM H_2_O_2_ as triggering stimulus (PT). The growth curves **(B)** and survival rates **(C)** were determined in naïve (C), primed (P), primed and triggered (PT) and triggered cells (T). The survival rates were calculated after 2 and 4 h of H O stress relative to untreated control cells. The results are from 3-4 biological replicates. Error bars represent the SD.

**Fig. S5. KatA and the peroxiredoxins AhpC, Tpx and Bcp are not involved in HOCl detoxification in *S. aureus*. (A-C)** Growth curves of *S. aureus* COL WT, Δ*katA* **(A)**, Δ*ahpC* **(B)**, Δ*tpx* **(C)** and Δ*bcp* mutants **(D)** in RPMI medium before (co) after exposure to 1.5 mM HOCl stress during the log phase. **(D)** Survival rates were determined as CFUs of *S. aureus* COL WT, Δ*katA*, Δ*ahpC*, Δ*tpx* and Δ*bcp* mutants at 2 and 4 h after treatment with 2.5 mM HOCl. Survival of the untreated control was set to 100%. Mean values and SD of 4-5 biological replicates are presented.

