## Supplemental Figures S1-S5 for "The catalase contributes to microaerophilic H_2_O_2_ priming and the peroxiredoxins AhpC, Tpx and Bcp confer resistance to organic hydroperoxides in *Staphylococcus aureus*"

Fig. S1

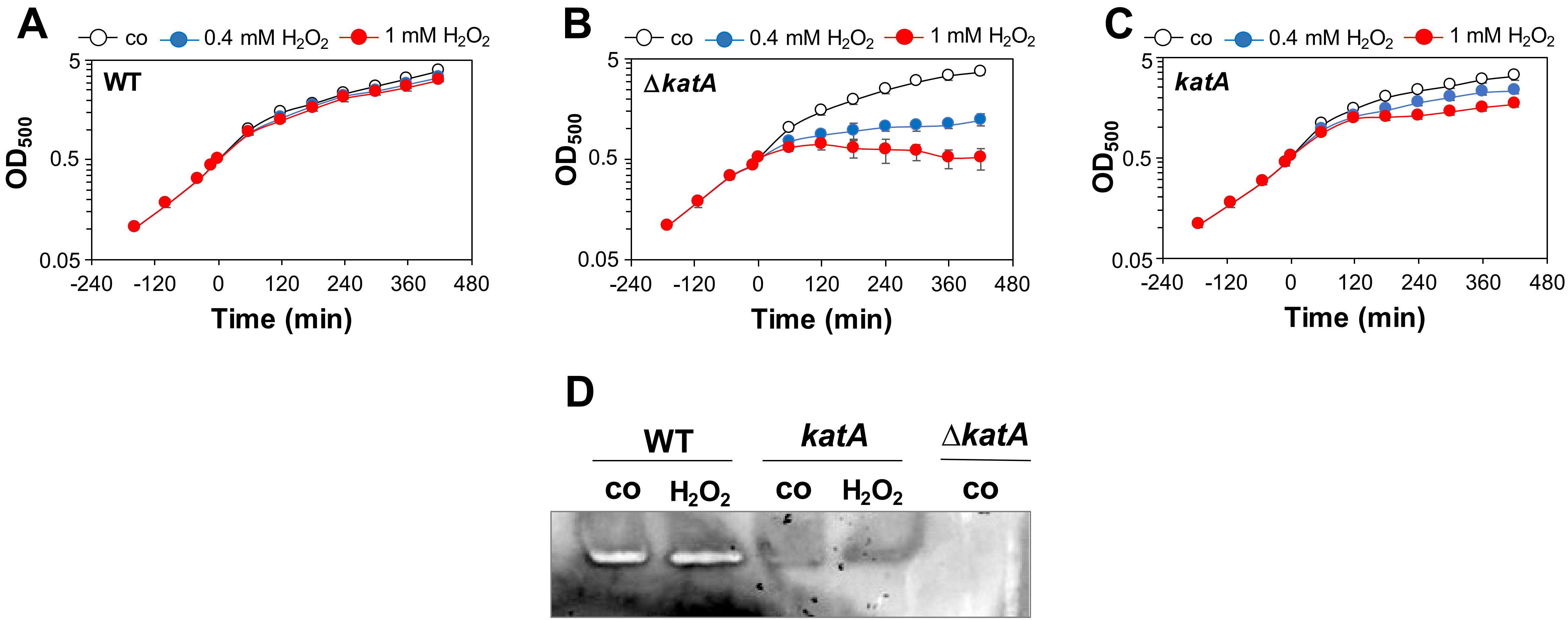


**Fig. S1. Complementation of KatA increases the $H_2O_2$ resistance in *S. aureus* (A-C), but the catalase activity in the *katA* complemented strain is lower compared to the WT (D).** Growth curves of the *S. aureus* COL WT **(A)**, the Δ*katA* mutant **(B)** and the *katA* complemented strain **(C)** after exposure to 0.4 and 1 mM $H_2O_2$ stress at an $OD_{500}$ of 0.5. Mean values and SD of four biological replicates are presented. **(D)** Catalase staining of aerobically grown *S. aureus* COL WT, the Δ*katA* mutant and the *katA* complemented strain before (co) and after exposure to 100 µM $H_2O_2$ at an $OD_{500}$ of 0.5. Protein extracts were separated by native PAGE and the catalase activity was determined using diamidobenzidine staining as described in the Methods.

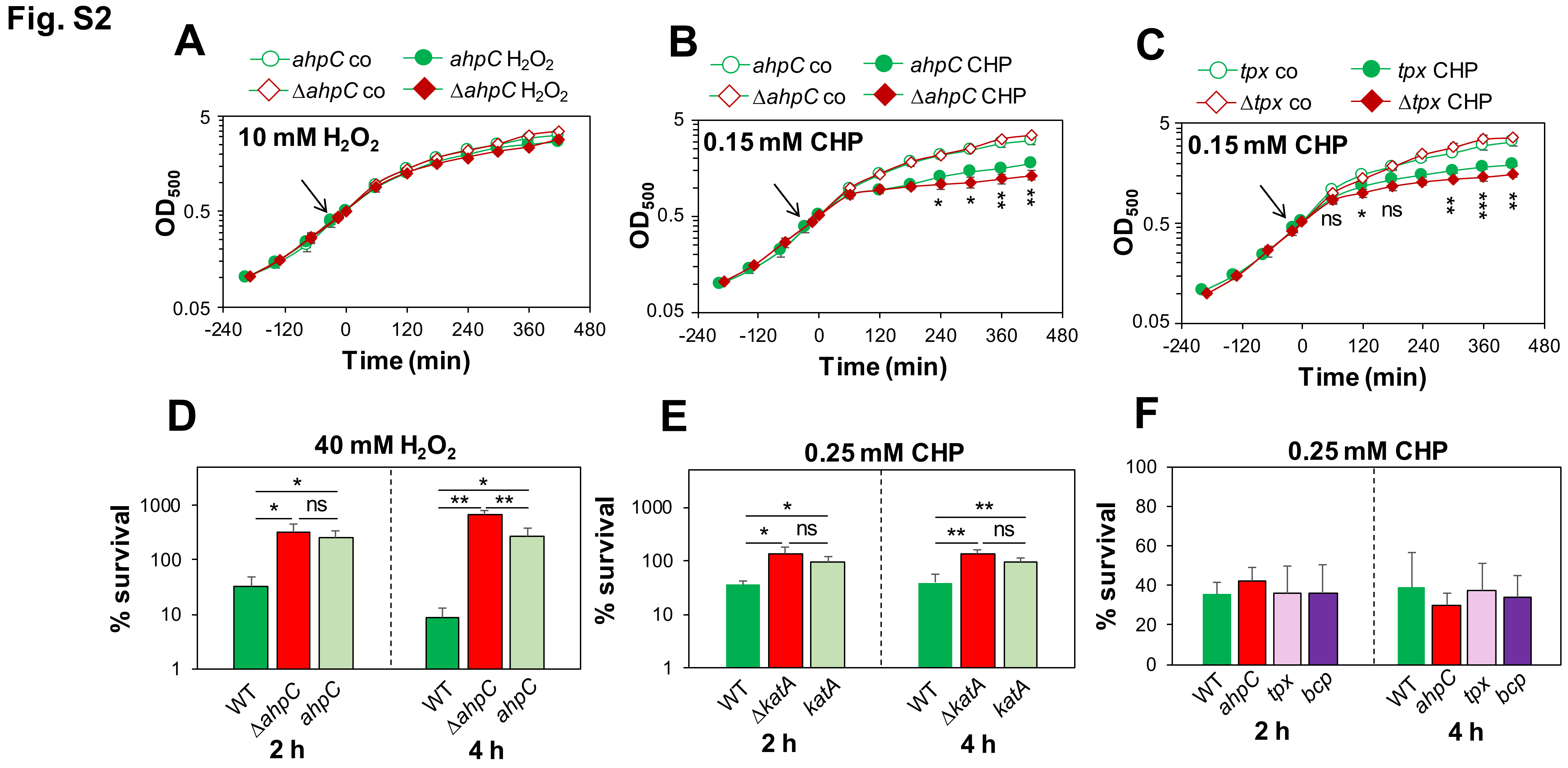


**Fig. S2. The CHP-sensitive phenotypes of the Δ*ahpC*, Δ*tpx* and Δ*bcp* mutants are abolished in the complemented strains. (A-C)** Growth curves of *S. aureus* COL WT, Δ*ahpC* and Δ*tpx* mutants and the complemented strains in RPMI medium before (co) and after exposure to 10 mM $H_2O_2$ **(A)** and 0.15 mM CHP **(B, C)**. **(D-F)** The survival rates were determined for the *S. aureus* COL WT, Δ*ahpC*, Δ*katA* and Δ*tpx* mutants and complemented strains after exposure to 40 mM $H_2O_2$ **(D)** and 0.25 mM CHP **(E, F)** based on CFU counts related to the untreated control, which was set to 100%. Mean values and SD of 4 biological replicates are presented. The statistics was determined using a Student's unpaired two-tailed t-test by graph prism. Symbols are: $^{ns}p > 0.05$, $^{*}p \leq 0.05$ ,$^{**}p \leq 0.01$ and $^{***}p \leq 0.001$.

**Fig. S3**

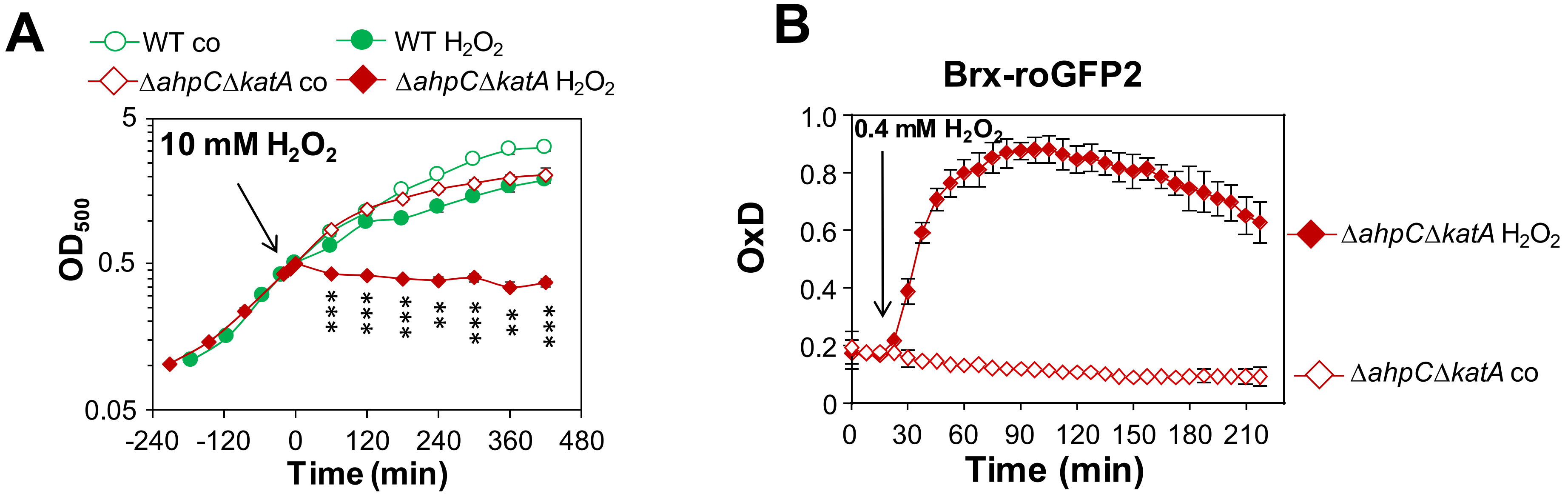


**Fig. S3. The Δ*katA ahpC* double mutant is hypersensitive towards $H_2O_2$ and delayed in recovery of the $E_{BSH}$ after 0.4 mM $H_2O_2$ (A)** Growth curves of *S. aureus* COL WT and the Δ*ahpC*Δ*katA* double mutant in RPMI medium before (co) and after exposure to 10 mM $H_2O_2$ at an $OD_{500}$ of 0.5. The survival rates were calculated relative to the untreated control, which was set to 100%. Mean values and SD of 3-4 biological replicates are shown. The statistics was calculated using the Student's unpaired two-tailed t-test by the graph prism software. Symbols are *p ≤ 0.05, **p ≤ 0.01 and ***p ≤ 0.001. **(B)** The Brx-roGFP2 biosensor in the *S. aureus* COL Δ*ahpC*Δ*katA* double mutant showed a fast oxidation and a delay in recovery of the reduced $E_{BSH}$ after 0.4 mM $H_2O_2$. The oxidation degree (OxD) of the Brx-roGFP2 response was calculated based on the 405/488 nm excitation ratios and normalized to fully reduced and oxidized controls. Mean values and SD of the OxD values are presented from 3 biological replicates.

**Fig. S4**

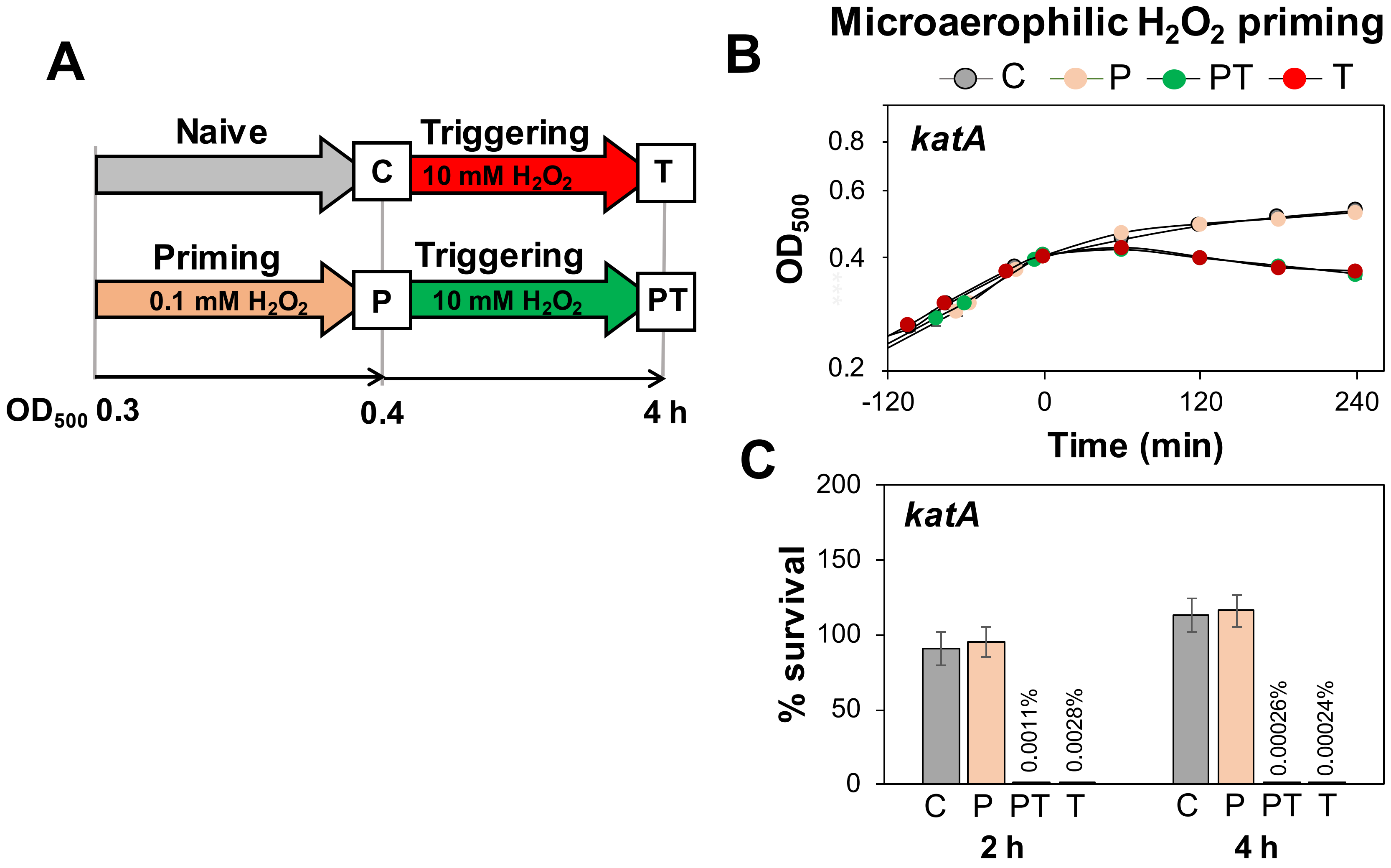


**Fig. S4. The *katA* complemented strain is not primable towards increased $H_2O_2$ resistance during the microaerophilic growth. (A)** Setup for microaerophilic priming and triggering experiments. The *S. aureus katA* complemented strain was primed during the log phase with 0.1 mM $H_2O_2$ for ~30 min **(P)** and subsequently treated with 10 mM $H_2O_2$ as triggering stimulus **(PT).** The growth curves **(B)** and survival rates **(C)** were determined in naïve (C), primed (P), primed and triggered (PT) and triggered cells (T). The survival rates were calculated after 2 and 4 h of $H_2O_2$ stress relative to untreated control cells. The results are from 3-4 biological replicates. Error bars represent the SD.

**Fig. S5**

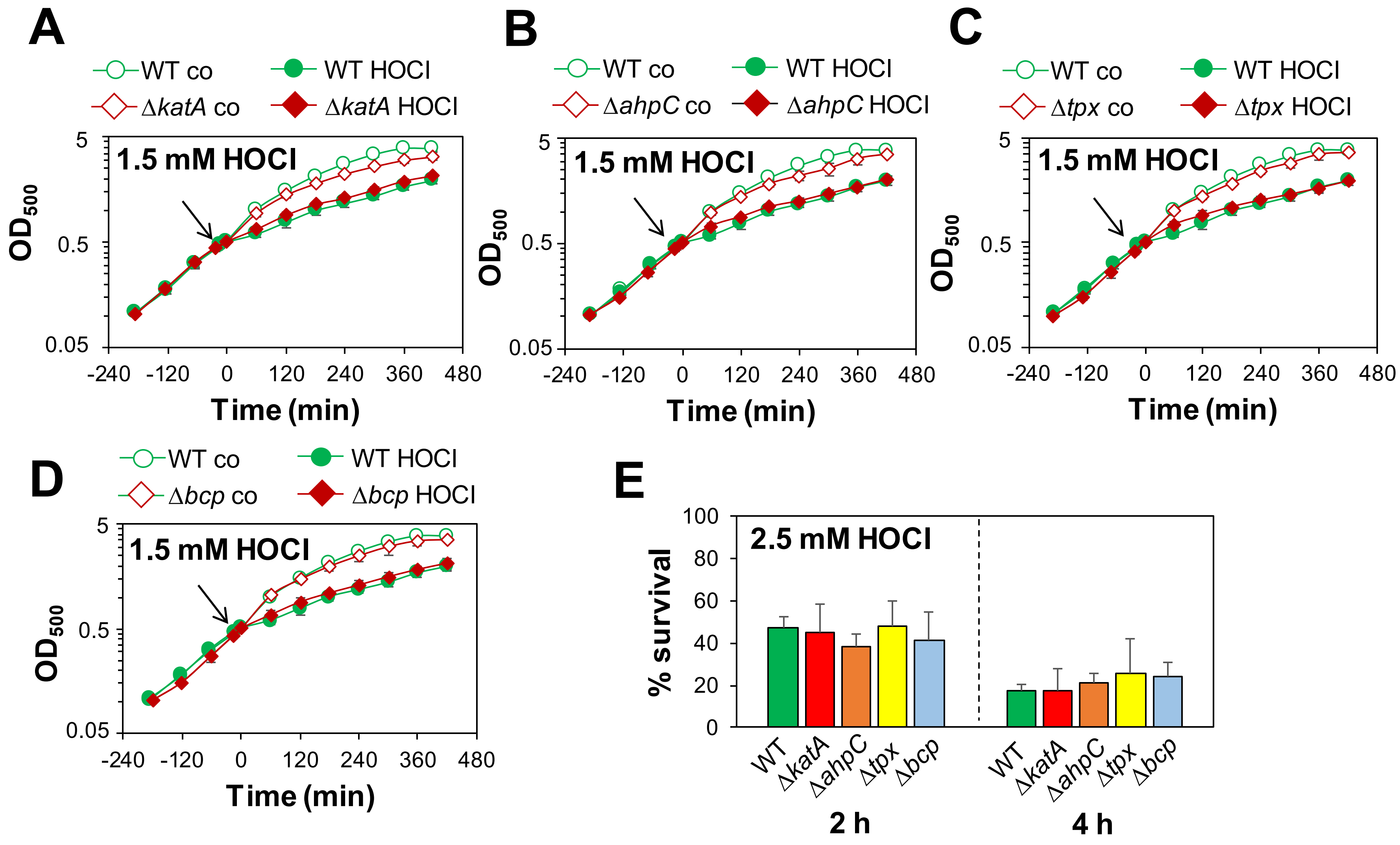


**Fig. S5. KatA and the peroxiredoxins AhpC, Tpx and Bcp are not involved in HOCl detoxification in *S. aureus*. (A-C)** Growth curves of *S. aureus* COL WT, Δ*katA* **(A)**, Δ*ahpC* **(B)**, Δ*tpx* **(C)** and Δ*bcp* mutants **(D)** in RPMI medium before (co) and after exposure to 1.5 mM HOCl stress during the log phase. **(D)** Survival rates were determined as CFUs of *S. aureus* COL WT, Δ*katA,* Δ*ahpC,* Δ*tpx* and Δ*bcp* mutants at 2 and 4 h after treatment with 2.5 mM HOCl. Survival of the untreated control was set to 100%. Mean values and SD of 4-5 biological replicates are presented.
