## Supplemental Tables S1-S3 for "The catalase contributes to microaerophilic H_2_O_2_ priming and the peroxiredoxins AhpC, Tpx and Bcp confer resistance to organic hydroperoxides in *Staphylococcus aureus*"

**Table S1. Bacterial strains**

| Bacterial strains | Description | Reference |
|---|---|---|
| ***Escherichia coli*** | | |
| DH5α | F-φ80dlacZ Δ(lacZYA-argF) U169 deoRsupE44ΔlacU169 (f80lacZDM15) hsdR17 recA1 endA1 (rk- mk+) supE44gyrA96 thi-1 gyrA69 relA1 | [1] |
| ***Staphylococcus aureus*** | | |
| RN4220 | restriction negative strain/MSSA cloning intermediate derived from 8325-4 | [2] |
| COL | Archaic HA-MRSA strain | [3] |
| COL-Δ*katA* | COL *katA* deletion mutant | [4] |
| COL-Δ*ahpC* | COL *ahpC* deletion mutant | This study |
| COL-Δ*bcp* | COL *bcp* deletion mutant | This study |
| COL-Δ*tpx* | COL *tpx* deletion mutant | This study |
| COL-Δ*perR* | COL *perR* deletion mutant | This study |
| COL-Δ*ahpC*Δ*katA* | COL *ahpC* and *katA* double deletion mutant | This study |
| COL pRB473-*brx-roGFP2* | COL WT expressing Brx-roGFP2 biosensor | [5] |
| COL-Δ*katA* pRB473-*brx-roGFP2* | COL *katA* deletion mutant expressing Brx-roGFP2 | This study |
| COL-Δ*ahpC* pRB473-*brx-roGFP2* | COL *ahpC* deletion mutant expressing Brx-roGFP2 | This study |
| COL-Δ*bcp* pRB473-*brx-roGFP2* | COL *bcp* deletion mutant expressing Brx-roGFP2 | This study |
| COL-Δ*tpx* pRB473-*brx-roGFP2* | COL *tpx* deletion mutant expressing Brx-roGFP2 | This study |
| COL-Δ*katA::*pRB473-*katA* | COL *katA* mutant complemented with pRB473-*katA* | [4] |
| COL-Δ*ahpC::*pRB473-*ahpC-His* | COL *ahpC* mutant complemented with pRB473-*ahpC-His* | This study |
| COL-Δ*bcp*::pRB473-*bcp-His* | COL *bcp* mutant complemented with pRB473-*bcp-His* | This study |
| COL-Δ*tpx::*pRB473-*tpx-His* | COL *tpx* mutant complemented with pRB473-*tpx-His* | This study |
| *Staphylococcus* phage 81 | | [6] |

**Table S2. Plasmids**

| Plasmid | Description | Reference |
|---|---|---|
| pRB473-Xyl | pRB373-derivative, *E. coli/ S. aureus* shuttle vector, containing xylose-inducible $P_{Xyl}$ promoter, $Amp^r$, $Cm^r$ | [7, 8] |
| pRB473-Xyl-*brx-roGFP2* | pRB473-derivative expressing *brx-roGFP2* under $P_{Xyl}$ | [5] |
| pRB473-Xyl-*katA* | pRB473-derivative expressing *katA* under $P_{Xyl}$ | [4] |
| pRB473-Xyl-*ahpC-His* | pRB473-derivative expressing His-tagged *ahpC* under $P_{Xyl}$ | This study |
| pRB473-Xyl-*bcp-His* | pRB473-derivative expressing His-tagged *bcp* under $P_{Xyl}$ | This study |
| pRB473-Xyl-*tpx-His* | pRB473-derivative expressing His-tagged *tpx* under $P_{Xyl}$ | This study |

**Table S3. Oligonucleotide primers**

| Primer name | Sequence (5' to 3') |
|---|---|
| pMAD-ahpC-for-BglII | CGCAGATCTCCACTCCTCGATACTTTACAAT |
| pMAD-ahpC-f1-rev | CTACTAAATCTAAACCAGGTTGCTGTAAATGGTAAGATTTCTTTG |
| pMAD-ahpC-f2-for | CAAAGAAATCTTACCATTTACAGCAACCTGGTTTAGATTTAGTAG |
| pMAD-ahpC-rev-SalI | CCAGTCGACCATCAATCATAGAATGCGTGAT |
| pMAD-bcp-for-BglII | CGCAGATCTCTTTTAGTATATGCACGTGCAA |
| pMAD-bcp-f1-rev | TGTTTTTAAGTTCTTCTATTTGTGTGGAAATTGTTCTCCTTTTTGC |
| pMAD-bcp-f2-for | GCAAAAAGGAGAACAATTTCCACACAAATAGAAGAACTTAAAAACA |
| pMAD-bcp-rev-SalI | CCAGTCGACGCTAACTTCGCAGTTCTAGTA |
| pMAD-tpx-for-BglII | CGCAGATCTACGCACGTTTACTCAATTTACA |
| pMAD-tpx-f1-rev | TTAAATATTTTTGTATGCAGCTAAACCTTTGAATGTTATTTCAGTCAT |
| pMAD-tpx-f2-for | ATGACTGAAATAACATTCAAAGGTTTAGCTGCATACAAAAATATTTAA |
| pMAD-tpx-rev-SalI | CCAGTCGACCAGGTAACACTTCTTTTACAGT |
| pMAD-perR-for-BglII | CGCAGATCTAATCACTTGAAAGCACATTACCA |
| pMAD-perR-f1-rev | TCTTGGCATTCTTTACAAACTCATTGATTCTATTTCAACACTCAT |
| pMAD-perR-f2-for | ATGAGTGTTGAAATAGAATCAATGAGTTTGTAAAGAATGCCAAGA |
| pMAD-perR-rev-SalI | CCAGTCGACGAATTTCAATAGTCAAATTTACAC |
| pRB-ahpC-for-BamHI- | TAGGGATCCATTCTTAGGAGGAAGATATTTATG |
| pRB-ahpC-rev-KpnI-His | CTCGGTACCTTAGTGATGGTGATGGTGATGGATTTTACCTACTAAATCTAAA |
| pRB-bcp-for-BamHI | TAGGGATCCAATGAAGAAAAAGGTGATTATATG |
| pRB-bcp-rev-KpnI-His | CTCGGTACCTCAGTGATGGTGATGGTGATGCCCCAAAATGTTTTTAAGTTCTT |
| pRB-tpx-for-BamHI | TAGGGATCCATGCAGGAGGTCAAGTATATGACTGAAATAACATTCAAAG |
| pRB-tpx-rev-SacI-His | CTCGAGCTCTTAGTGATGGTGATGGTGATGAATATTTTTGTATGCAGC |
| katA-NB-for | AAAGGTTCTGGTGCATTTGG |
| katA-NB-rev | CTAATACGACTCACTATAGGGAGAAATGTGTTCCTCCACCTTGG |
| ahpC-NB-for | TCCTGCTGACTTCTCATTCGT |
| ahpC-NB-rev | CTAATACGACTCACTATAGGGAGAGGTTGCAATGTTTTAGCGCC |

Restriction sites are underlined.
